# Gradual opening of Smc arms in prokaryotic condensin

**DOI:** 10.1101/2021.01.21.427566

**Authors:** Roberto Vazquez Nunez, Yevhen Polyhach, Young-Min Soh, Gunnar Jeschke, Stephan Gruber

**Author notes:** Equal contribution (Roberto Vazquez Nunez, Yevhen Polyhach). Corresponding author, Lead Contact, E-mail: S.G.

## Abstract

Multi-subunit SMC ATPases control chromosome superstructure apparently by catalyzing a DNA-loop-extrusion reaction. SMC proteins harbor an ABC-type ATPase ‘head’ and a ‘hinge’ dimerization domain connected by a coiled coil ‘arm’. Two arms in a SMC dimer can co-align, thereby forming a rod-shaped particle. Upon ATP binding, SMC heads engage, and arms are thought to separate. Here, we studied the shape of *B. subtilis* Smc-ScpAB by electron-spin resonance spectroscopy. Arm separation was readily detected proximal to the heads in the absence of ligands, while separation near the hinge largely depended on ATP and DNA. Artificial blockage of arm opening eliminated DNA stimulation of ATP hydrolysis, but did not prevent basal ATPase activity. We identified an arm-to-arm contact as being important for controlling the molecular transformations. Point mutations at this arm interface eliminate Smc function. We propose that partially open, intermediary conformations provide directionality to SMC DNA translocation by binding suitable DNA substrates.

## Introduction

SMC protein complexes are ancient enzymes with a unique architecture that organize chromosomal DNA molecules, presumably by DNA loop extrusion (Yatskevich et al., 2019). In eukaryotes, cohesin folds DNA into loop domains to regulate gene expression and to direct DNA recombination (Ba et al., 2020; Merkenschlager and Nora, 2016). By a distinct mechanism, cohesin also holds sister chromatids together (Yatskevich et al., 2019). In mitosis, condensin folds DNA into a series of loops that are dynamically anchored along a chromatid axis thus supporting chromosome condensation and sister chromatid resolution (Earnshaw and Laemmli, 1983; Gibcus et al., 2018; Marsden and Laemmli, 1979; Naumova et al., 2013). The essential functions of another relative, the Smc5/6 complex, are less well understood (Aragón, 2018).

In many bacteria, Smc-ScpAB complexes initiate a loop-extrusion-type activity at one or few selected starting points that are defined by 16 bp *parS* DNA sequences. *parS* sites are located in the replication origin region. They recruit the clamp-like CTP-binding protein ParB (Jalal and Le, 2020; Osorio-Valeriano et al., 2019; Soh et al., 2019), which in turn promotes the loading of Smc-ScpAB complexes onto the chromosome (Gruber and Errington, 2009; Minnen et al., 2016; Sullivan et al., 2009; Wilhelm et al., 2015). Bidirectional translocation of Smc-ScpAB away from a *parS* site brings together the flanking DNA sequences, thus co-aligning the left and the right arm of the chromosome (Minnen et al., 2016; Tran et al., 2017; Wang et al., 2017; Wang et al., 2015). This translocation is thought to localize DNA entanglements on the replicating chromosome (*i*.*e*. knots and catenanes) facilitating chromosome individualization by DNA topoisomerase (Bürmann and Gruber, 2015; Gruber et al., 2014; Orlandini et al., 2019; Wang et al., 2014).

Smc-ScpAB complexes translocate rapidly (∼1 kb/sec) along chromosomal DNA *in vivo. In vitro* they support only limited ATP hydrolysis activity (< 1/sec), implying a large motor step size (∼1 kb or ∼600 nm) (Hirano and Hirano, 2006; Vazquez Nunez et al., 2019; Wang et al., 2017; Wang et al., 2018). Several models for Smc translocation have been proposed (Diebold-Durand et al., 2017; Hassler et al., 2018; Marko et al., 2019; Terakawa et al., 2017). Smc proteins are comprised of an ABC-type ATP-binding head domain and a hinge domain connected at a distance by a long antiparallel coiled coil arm. Hinge domains form homotypic interactions in prokaryotic Smc complexes (Haering et al., 2002). Smc dimers furthermore associate with a kleisin subunit, in bacteria named ScpA. Via its amino- and carboxy-terminal domains, ScpA bridges the head of one Smc protein with the head-proximal arm of the other (Bürmann et al., 2013). This generates tripartite SMC-kleisin rings that entrap chromosomal DNA double helices (Gligoris et al., 2014; Wilhelm et al., 2015). The kite subunit ScpB also forms dimers that associate with the central region of ScpA (Bürmann et al., 2013).

Smc arms contact one another. They co-align lengthwise thus collapsing the Smc-ScpAB complex into a rod-shaped particle (Diebold-Durand et al., 2017; Minnen et al., 2016; Soh et al., 2015). Eight distinct contacts are found between the two Smc arms, four of which involve amino-terminal sequences (1^N^, 4^N^, 6^N^, and 8^N^; see Figure 2B) and the other four carboxy-terminal sequences (2^C^, 3^C^, 5^C^, and 7^C^) (Diebold-Durand et al., 2017). In yeast condensin, corresponding sequences are also found in juxtaposition (Lee et al., 2020) (Figure S4D). In some SMC complexes (including cohesin and condensin), the arms fold at an ‘elbow’, thus bringing the hinge into proximity of the heads (Bürmann et al., 2019). Such folding has not yet been observed for bacterial Smc and its role is unclear. Smc heads engage with one another by sandwiching two ATP molecules using active site residues provided by both heads thereby forming the catalytic center for ATP hydrolysis (Hirano et al., 2001; Hopfner, 2016; Lammens et al., 2004). ATP-engagement of Smc heads is thought to be incompatible with full arm alignment (Diebold-Durand et al., 2017; Kamada et al., 2017; Lammens et al., 2004; Muir et al., 2020) thus delineating two mutually exclusive conformations, one with ATP-engaged heads (O-shaped ‘open’ conformation) and one with completely aligned arms (I-shaped ‘closed’ conformation). In the open conformation, ATP-engaged heads divide the lumen of the SMC-kleisin ring into the S compartment that is encircled by the long arms and the hinge, and the K compartment which is enclosed by ScpAB. The open conformation of *B. subtilis* (*Bsu*) Smc-ScpAB harbors two sites for DNA binding in the S compartment: a hinge/DNA interface and a head/DNA interface (Vazquez Nunez et al., 2019). In the closed conformation, the Smc arms are aligned. The S compartment is thus closed and separated from the K compartment by juxtaposed Smc heads. Principally consistent observations have recently been made by cryogenic electron microscopy for other SMC complexes, whereas AFM studies indicated much higher flexibility in the SMC arms (see Discussion). SMC arm dynamics appear crucial for DNA loop extrusion, but exactly how the different conformations bind to DNA and contribute to DNA translocation remains to be elucidated.

Here, we studied the shape of the *Bsu* Smc-ScpAB by EPR spectroscopy. We observed in addition to the open and the closed conformation an intermediary one with partially open S compartments. Separation of arms was detected near the heads even in the absence of ligands. Opening at the hinge, however, is largely dependent on ATP and DNA binding. We showed that partial opening of the S compartment is sufficient for ATP hydrolysis, but intriguingly DNA stimulation of the ATPase activity requires complete opening. This implies that an open S compartment is an integral part of the normal ATP hydrolysis cycle. We identified one out of the eight arm contacts as particularly important for controlling the dynamic Smc architecture. Mutating a single residue at this 4^N^ contact interface eliminates Smc function, presumably by preventing arm closure. The mutations cause defect in chromosome loading or DNA translocation. Altogether, our results suggest that the S compartment opens by a graded transition starting at the heads. Head-proximal arm contacts dissociate relatively easily, while hinge-proximal contacts are more stably engaged. We identified the 4^N^ contact as critical for controlling the opening/closure reaction. It may provide directionality to the DNA translocation motor.

## Results

### Conformations of Smc-ScpAB detected by EPR

Here, we aimed to characterize the conformational ensemble of Smc-ScpAB by measuring arm-to-arm distances. Based on published work, we expected the distances to be broadly distributed covering a spectrum of closed and open conformations. Such dynamic structures are difficult to assess by high resolution techniques such as X-ray crystallography and electron microscopy. Therefore, we employed electron paramagnetic resonance (EPR-DEER) which detects the coupling of electron spins over a larger distance range (from 1.5 to 10 nm) and yields information on distance distributions for a population of particles (Jeschke, 2012; Polyhach et al., 2012; Reginsson and Schiemann, 2011). We labelled purified cysteine-mutant Smc protein at one out of four selected positions (D193C, E217C, R643C and R718C) with the spin label MTSL and reconstituted holo-complexes by mixing with unmodified ScpA and ScpB (Figure 1A). The EPR experiments were performed with the cysteine-lite Smc^3S^ protein (C119S, C437S and C826S) to minimize any off-target labelling. MTSL-labelling efficiencies were around 90 % as calculated from continuous-wave EPR spectra (Figure S1A). Of note, protein samples were prepared in deuterated environment to reduce electron spin relaxation induced by proton nuclear spin diffusion (El Mkami and Norman, 2015). Deuterated buffers had only minor impact on protein function, as judged by near-normal ATPase activity (Figure 1SB). For most experiments, the Smc protein also included the Walker B E1118Q (EQ) mutation which hinders the ATP hydrolysis step. Protein preparations were pre-incubated with or without ATP and linear 40 bp double-stranded DNA, designated as dsDNA_40_, flash-frozen in liquid nitrogen and measured at a temperature of 50 K.

**Figure 1.**
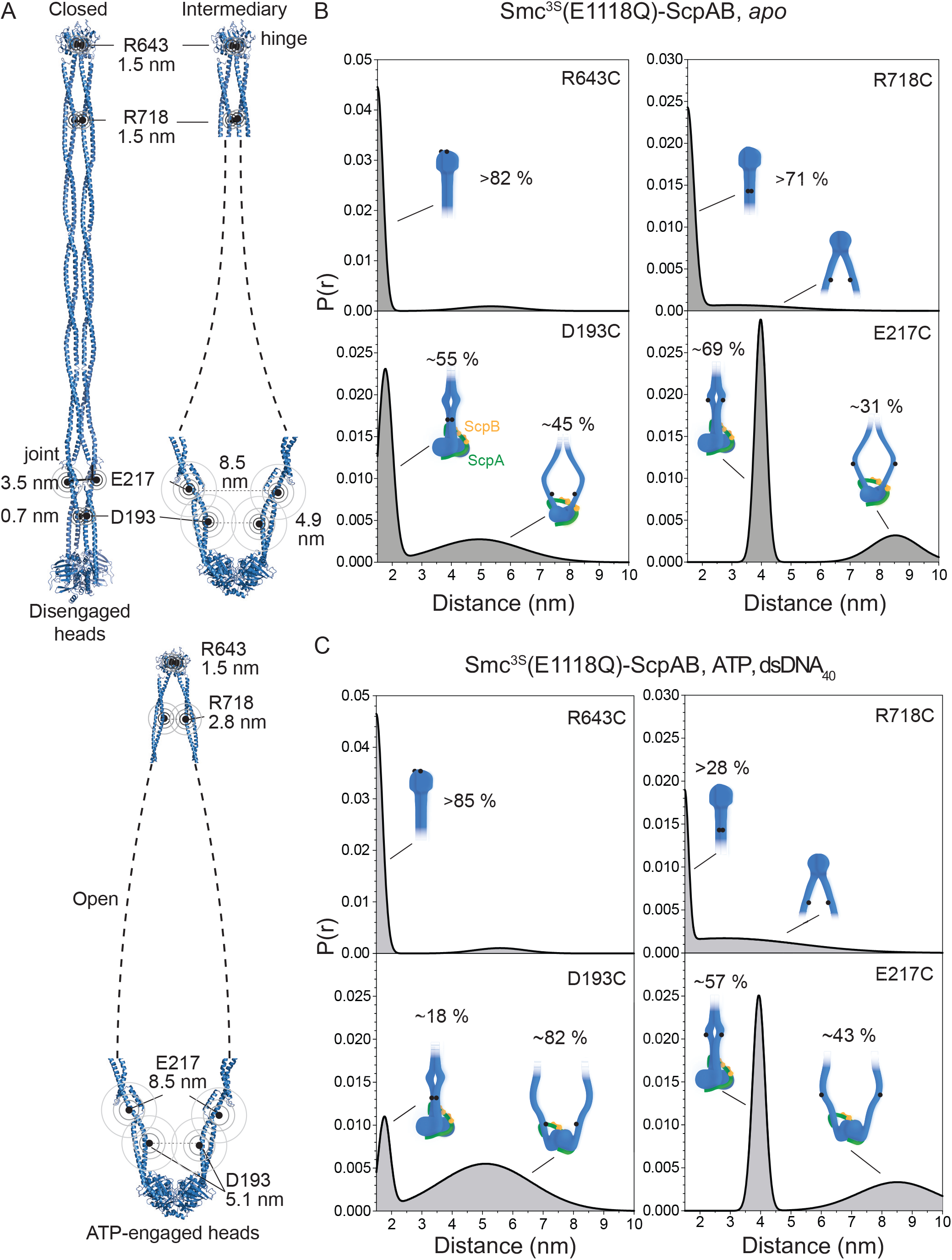
Distance distributions in Smc-ScpAB determined by electron spin resonance (EPR-DEER). A. Structural models of Smc dimers in the closed (top left panel), an intermediary (top right panel), and the open conformation (bottom panel). Models were built with the help of available crystal structures (Diebold-Durand et al., 2017; Soh et al., 2015). Dashed lines indicate putative bending of the Smc arms. Concentric circles indicate the positions of the cysteines used for MTSL labelling. Distances are maxima for sub-populations as observed by EPR. B. Probability distributions as measured by EPR-DEER with reconstituted Smc-ScpAB in the absence of ligands (apo). Schematics indicate the conformation for sub-populations with estimated relative abundance given. C. Probability distributions of Smc-ScpAB measured in the presence of 1 mM ATP and 10 µM double-stranded 40 bp DNA (dsDNA_40_). Display as in (B).

Most distances measured across the hinge domain dimer in Smc^3S^(R643C, E1118Q) were narrowly distributed with a peak at ∼1.5 nm, at the lower edge of the sensitive range of high-power Q-band DEER, well-fitting with available crystal structures (Cα-Cα ∼1.1 nm) (Figures 1A and 1B) (Griese and Hopfner, 2011; Haering et al., 2002; Kamada et al., 2017; Soh et al., 2015). A minor fraction with larger distances (∼4-6 nm) was potentially also noticeable (Figure 1B). Arm-to-arm distances near the heads showed a clear bimodal distribution in the absence of ligand (the *apo* state) (Figure 1B, S1C, S1E-F). For Smc^3S^(D193C, E1118Q) (at contact 1^N^), a population with narrowly distributed short distances represented about 55 % of all distances. These were centered at ∼1.7 nm and displayed a good fit with the Smc rod model built from crystal structures (Diebold-Durand et al., 2017). The long-distance population showed a much broader distribution spanning from about 3.0 to at least 6.0 nm, indicating separated Smc arms near the heads in a substantial proportion of Smc complexes (Figure 1B). A similar pattern was observed with Smc^3S^(E217C, E1118Q) (at contact 2^C^): ∼69 % of the dipolar couplings resided in a narrow short-distance population centered at ∼3.9 nm, while other distances ranged from ∼6.0 nm to more than 8.5 nm (Figure 1B). Again, the short-distance population is in good agreement with the rod model (Diebold-Durand et al., 2017). For the hinge-proximal arm position, Smc^3S^(R718C, E1118Q) (at contact 7^C^), 71 % of the distances accumulated in a population with short arm-to-arm distances at ∼1.5 nm. This number likely underestimates the size of the population due to partial suppression of the dipolar modulation at such short distances (similar to R643C) (Figure 1B). These measurements imply that in the absence of ligands one or more conformations exist in addition to the closed rod structure. Smc arms appeared separated near the heads (especially at contact 1^N^) in a larger fraction of complexes than near the hinge, implying the presence of partially open, intermediary conformations (Figure 1A). This is consistent with the pattern of arm cross-linking observed *in vivo*, where the levels of cysteine cross-linking were somewhat higher at contacts 8^N^ and 6^N^ than at contacts 3^C^, 2^N^ and 1^C^ (Diebold-Durand et al., 2017).

Addition of ATP to Smc(E1118Q)-ScpAB did not noticeably change the distribution between the short- and the long-distance population for S217C and R643C (Figure S1D). For D193C, the fraction of long distances became slightly larger indicating a trend towards arm opening upon ATP binding. For S217C, the long distances became somewhat shorter. The distance distribution at the hinge (R643C) remained virtually unchanged also in the presence of ATP and dsDNA_40_ (Figure 1D). Arm-to-arm distances, however, showed a pronounced shift from the short-distance to the now predominant long-distance population upon pre-incubation with ATP and dsDNA_40_. For example, close to the hinge (R718C) (at contact 7^C^), a broad long-distance population ranging from about 2.0 to 7.0 nm represented about 70 % of all measurable distances in the presence of ATP and dsDNA_40_ (Figure 1C). The extent of shift to larger distances varied slightly between the cysteine positions, possibly resulting from uncertainty in quantifying short-distance (<1.5 nm) or long-distance (>10 nm) populations or indicating that the cysteine residues or their chemical labelling mildly affected the stability of the conformations. Regardless, these results provided strong support for the notion that Smc arms separate from one another upon ATP and DNA addition. The Smc arms are presumably fully detached in a significant fraction of complexes when bound to ATP and DNA. From the distance distribution at R718C, we estimated that the arms are connected to the hinge at narrow angles (0 to ∼45°). More open angles—as seen in some crystal structures of Smc hinge fragments (∼180°)—were however not observed. The open conformation thus represents elongated, oval-shaped particles. Of note, we obtained similar trends in the absence of ScpAB and when using wild-type Smc instead of Smc(E1118Q) proteins (Figure S1E-F), although arm dissociation (at D193C) with ATP and DNA was less pronounced with wt Smc when compared to Smc(E1118Q).

Altogether, we conclude from the EPR measurements that *apo* Smc-ScpAB exists mainly in the closed conformation and in smaller sub-populations of the open and very likely also of an intermediary conformation. Upon ligand binding, the distributions shift towards larger arm-to-arm distances generating a sizeable fraction of open conformations. The broad range of measured distances indicates that the arms are somewhat flexible at least when arm alignment is lost.

### Partial arm opening is sufficient for ATP hydrolysis, but DNA stimulation requires full opening

We next wondered whether intermediary conformations—as observed by EPR–are able to support ATP hydrolysis or whether a fully open conformation is prerequisite for ATP hydrolysis. To test this, we blocked S compartment opening by engineering a covalent arm-to-arm junction at selected arm positions: at contacts 1^N^, 3^C^, 4^N^, and 7^C^ (Figure 2B). Stiff arms would be expected to prevent ATP hydrolysis when conjoined, while flexible arms would support normal ATP hydrolysis even when conjoined. A reduction in the ATPase rate varying with the position of the engineered arm-to-arm junction would imply limited flexibility.

**Figure 2.**
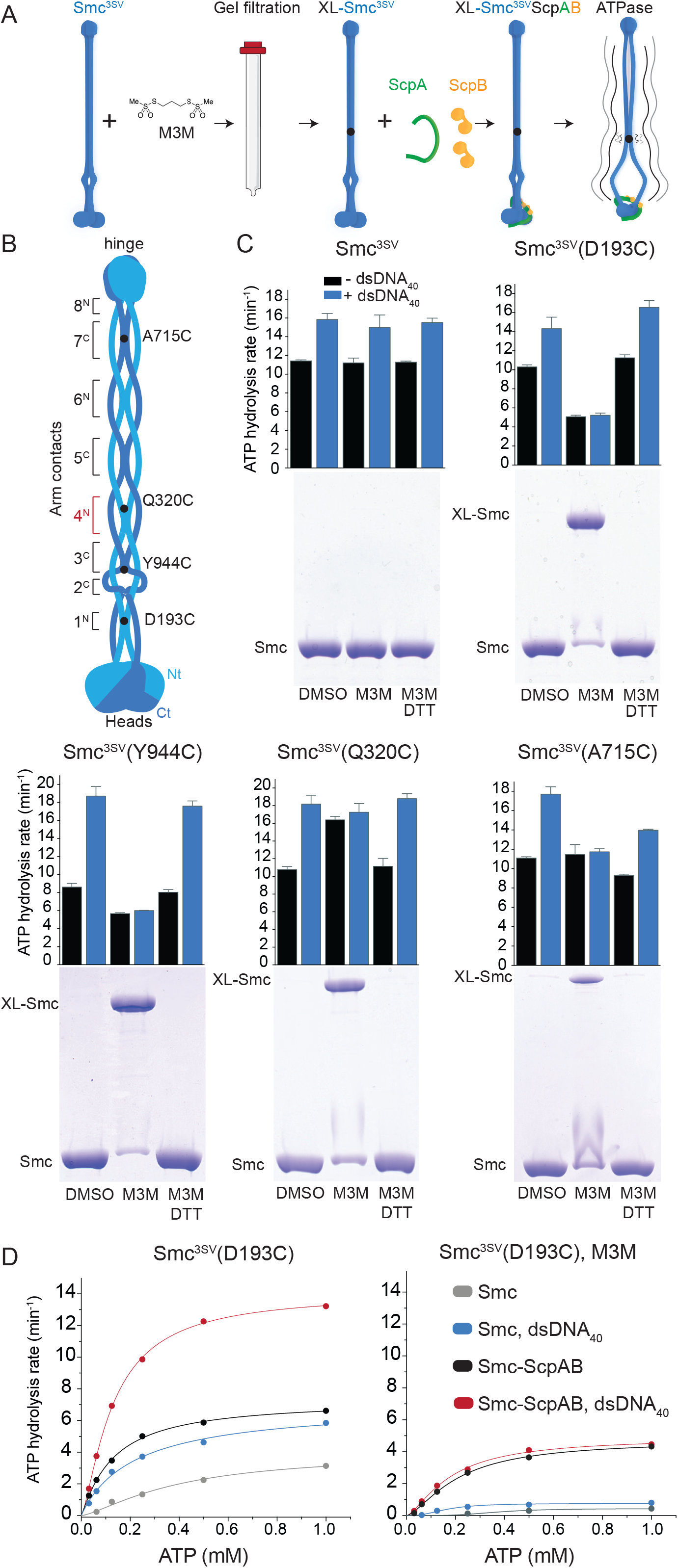
ATP hydrolysis by Smc-ScpAB with cross-linked arms. A. Schematic for the preparation of cross-linked protein for ATPase measurements. Single cysteine Smc^3SV^ proteins were purified and incubated with M3M cross-linker. Excess of M3M and any protein aggregates were removed by gel filtration. Eluate fractions were mixed with stoichiometric amounts of purified ScpA and ScpB to reconstitute holo-complexes. ATPase activity was determined. B. Positions of arm contacts and engineered cysteine residues in the *Bsu* Smc dimer in the closed conformation (Diebold-Durand et al., 2017). Amino-terminal sequences (Nt) are shown in light blue colours and carboxy-terminal sequences (Ct) in dark blue colours. Inter-subunit arm contacts are numbered 1 to 8 from the heads towards the hinge. Superscripts (N or C) indicate the involvement of amino- or carboxy-terminal sequences at the contact, respectively. Black circles denote the positions of cysteine residues engineered for site-specific crosslinking. C. ATP hydrolysis rates for preparations of Smc-ScpAB with cross-linked arms. Top panels: bar graphs of ATP hydrolysis rates per Smc protein measured with 1 mM ATP, 0.15 µM Smc-ScpAB complex in the absence (black bars) and presence (blue bars) of 3 µM dsDNA_40_. Error bars correspond to the standard deviation calculated from 3 technical replicates. Bottom panels: Coomassie-stained SDS-PAGE gel of cross-linked Smc protein. Solvent only (‘DMSO’), cross-linker only (‘M3M’), cross-linker with subsequent treatment with reducing agent (‘DTT’). D. ScpAB dependence of the ATP hydrolysis rates of cross-linked Smc^3SV^(D193C). Hydrolysis rates were measured at 0.15 µM protein concentration in the presence and absence of stoichiometric amounts of ScpAB and 3 µM dsDNA_40_. Non-crosslinked control (left panel), M3M cross-linked proteins (right panel). Lines represent non-linear regression fits to the Hill-model.

We again made use of site-directed chemical modification. We cross-linked cysteine residues in purified preparations of Smc with the thiol-reactive compound 1,3-propanediylbismethanethiosulfonate (M3M) to then reconstitute arm-linked holo-complexes and measure their ATPase activity (Figure 2A). M3M cross-linking is efficient as well as reversible thus allowing to break the engineered arm-to-arm connection to recover lost activity. These experiments were performed with cys-free Smc^3SV^ (C119S, C437S, C826S, C1114V) protein (Figure 2C). However, comparable results were obtained with the Smc^3S^ variant (Figure S2A). Incubation of Smc^3SV^(D193C), Smc^3SV^(Q320C), Smc^3SV^(A715C) or Smc^3SV^(Y944C) with M3M produced at least 70 % to 85 % of chemically cross-linked dimers as judged by non-reducing SDS-PAGE analysis. Addition of up to 10 mM DTT was required to reverse the cross-linking reaction (Figure 2C).

All cross-linked preparations yielded significant ATPase activity regardless of the position of the engineered arm-to-arm junction. While a Cys-free control sample showed no discernible effect of M3M treatment on ATP hydrolysis (Figure 2C), a substantial loss of ATPase activity was observed with head-proximal cysteine residues (D193C at contact 1^N^ and Y944C at contact 3^C^). No such effect was seen for the more distant positions (Q320C and A715C at contacts 4^N^ and 7^C^, respectively). In all cases, normal ATPase activity was restored by pre-incubation with reducing agent. These results strongly suggest that complete separation of Smc arms is not required for productive head engagement and for ATP hydrolysis. Intermediary conformations may support ATP hydrolysis. Cross-linking of Q320C (at contact 4^N^) even increased the basal ATPase rate (see below).

We found that the stimulation of the ATPase activity by addition of 40 bp double-stranded DNA (dsDNA_40_) was eliminated by M3M treatment regardless of the position of the cysteine residue on the arm and regardless of any dampening or stimulating effect of the arm-to-arm junction on ATP hydrolysis (Figure 2B). DNA stimulation was however restored upon pre-incubation with reducing agent. The stimulation of the Smc ATPase by dsDNA_40_ thus depends on the separation of residues on opposite arms, being consistent with the idea that complete arm opening is required for DNA stimulation. DNA binding at the hinge/DNA interface may boost ATP hydrolysis by modifying the organization of the arms as previously proposed (Soh et al., 2015). In contrast to the strong effect on ATP hydrolysis (Figure 2D), the arm-to-arm junction at Y944C did not substantially hinder DNA binding by Smc-ScpAB (Figure S2B), suggesting that ATP hydrolysis is uncoupled from DNA binding in the cross-linked material or that DNA binding predominantly occurs at a site this is uncoupled from the ATPase even in unmodified Smc-ScpAB.

Altogether, these findings imply that the Smc ATPase can operate in two modes: as a basal ATPase in a partially open conformation and as a DNA-stimulated ATPase presumably with an open conformation. The separation of head-proximal arms promotes ATP hydrolysis, while hinge-proximal arms may remain juxtaposed during basal ATP hydrolysis cycles. Interestingly, ScpAB becomes essential for ATP hydrolysis when arms are conjoined at contact 1^N^ (D193C) (Figure 2D), suggesting that it helps to overcome constraints imposed by the engineered junction on the ATPase heads. Of note, the fact that DNA is unable to stimulate ATP hydrolysis in arm-locked complexes is consistent with the prior observation that the hinge/DNA interface but not the head/DNA interface is important for DNA stimulation of ATP hydrolysis (Vazquez Nunez et al., 2019).

### Stimulation of ATP hydrolysis by chemical modification of Smc arms

During the above experiments, we noticed that another cysteine variant, Smc^3SV^(E295C), showed aberrant enzyme kinetics. The protein supported robust cysteine cross-linking (87.8 ± 4.6 %) with the cross-linked protein preparation displaying a ∼2.5-fold higher ATPase activity (Figure 3C, S3A and S3B). This increase was reversed to normal levels by pre-incubation with reducing agent. The E295C mutation largely eliminated DNA stimulation of the ATPase even without cross-linking. This finding confirmed that robust ATP hydrolysis can be achieved when arms are covalently conjoined (at contact 4^N^) and suggests that Smc arms can have a positive effect on ATP hydrolysis even when they are artificially linked together.

**Figure 3.**
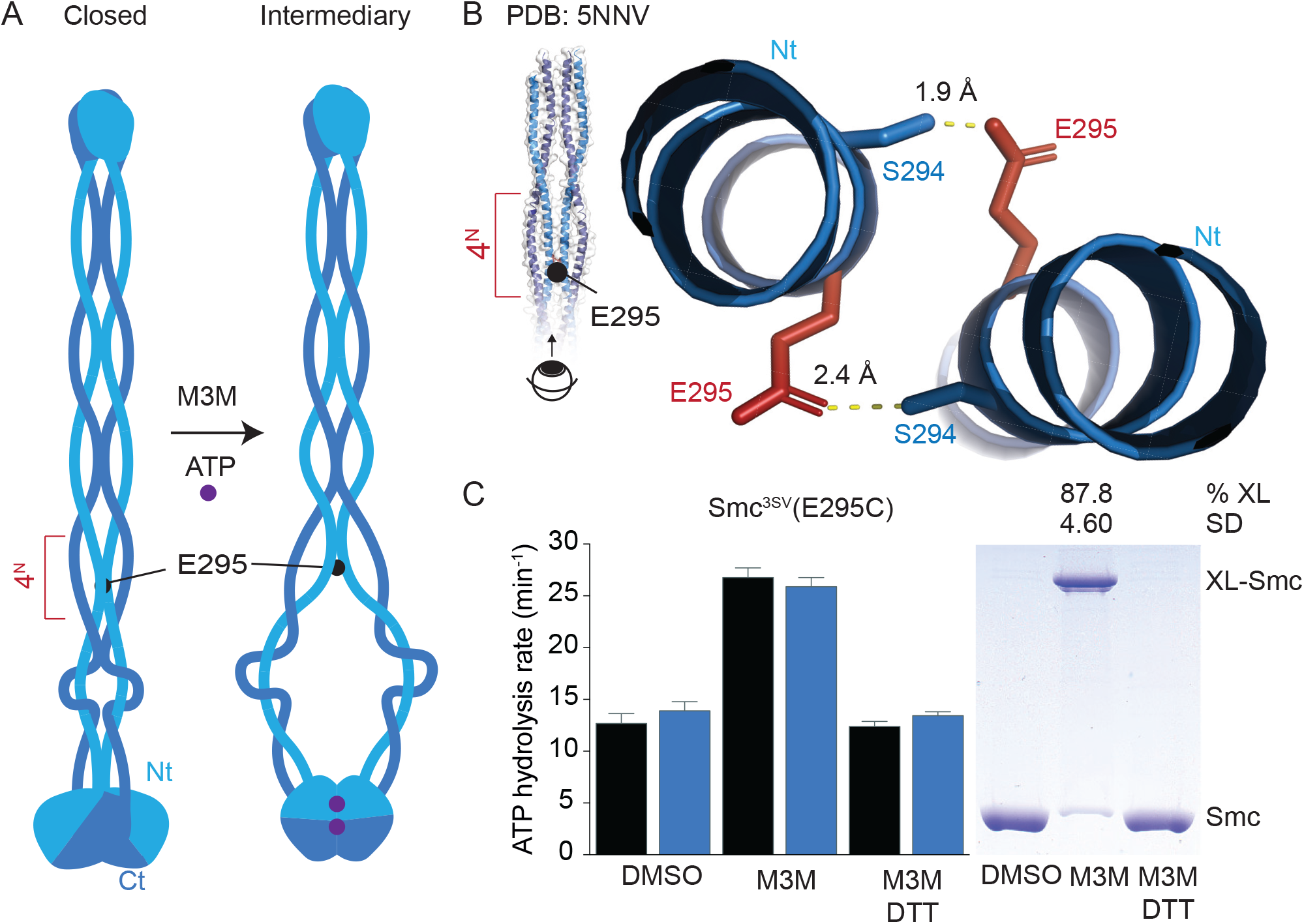
Cross-linking of E295C at the 4N arm contact enhances ATP hydrolysis. A. Schematic of arm contacts in the Smc dimer in closed and intermediary conformations. The position of E295 is indicated as black dot and the 4^N^ arm contact is denoted by a red bracket. ATP molecules are represented as purple dots. B. Structural organization of the 4^N^ arm contact in *Bsu* Smc (PDB: 5NNV) in cartoon representation. Hydrogen bonds are indicated as dotted lines in yellow colours. C. ATP hydrolysis rates and SDS-PAGE of cross linked Smc(E295C). Display as in Figure 2C.

In the crystal structure (PDB: 5NNV), E295 residues form inter-subunit hydrogen bonds with S294 residues (Figure 3B), thus likely stabilizing the 4^N^ arm contact (Diebold-Durand et al., 2017). The lack of these hydrogen bonds in the E295C mutant presumably interferes with normal arm alignment. Cross-linking of E295C may further disrupt Smc rod alignment. Intriguingly, we previously isolated point mutations at the 4^N^ arm contact (*i*.*e*. D280G, Q320R, or E323K) that suppressed the lethal phenotype of an arm length-modified Smc protein (Bürmann et al., 2017), supporting the notion that the 4^N^ arm contact is particularly important for controlling conformational transitions during the ATP hydrolysis cycle.

### A critical contact for arm alignment

To discern the physiological relevance of the 4^N^ arm contact, we next looked at the conservation of Smc arm sequences at this interface. Residue G302 and neighboring residues are well conserved in firmicute Smc proteins while arm sequences are otherwise relatively poorly conserved (Figure 4A). The glycine residue is located directly at the interface (Figure 4B), separated by two α-helical turns from E295. To test whether G302 is important for Smc function, we generated ten substitutions by allelic replacement at the endogenous *smc* locus in *B. subtilis* (Figure 4C). Remarkably, substitution to glutamate (GE), tryptophan (GW), or phenylalanine (GF) prevented growth of *Bsu* on nutrient rich medium—similar to *smc* deletion mutants— despite the GE and GW proteins being expressed at normal levels (Figure 4D). Three other mutants (GM, GQ, GK) were unable to support growth of a *ΔparB* mutant on this growth medium, while the arginine mutant (GR) supported growth of the double mutant only poorly (Figure S4A). Substitutions for residues with smaller side chains (alanine, valine and serine) had no discernible effects on growth. Thus, residues with negatively charged or bulky sidechains at position 302 hinder protein function, presumably by destabilizing the 4^N^ arm contact. To test this, we combined the G302 mutations with a sensor cysteine for arm alignment (A715C, at contact 7^C^) and one for head juxtaposition (S152C). In these experiments, we focused on two G302 mutants with a severe growth defect and sidechains with distinct physicochemical properties (GE and GW). The two mutations indeed reduced cysteine cross-linking at both sites *in vivo*, with GW having rather mild and GE quite dramatic effects (Figure 4E). The GW mutation also strongly reduced the abundance of the closed conformation (i.e. the short distance population) as measured by EPR with D193C in the absence and presence of ATP and DNA (Figure S4B, Table S2). We conclude that residue G302 and the 4^N^ arm contact are important for co-aligning arms in the closed conformation.

**Figure 4.**
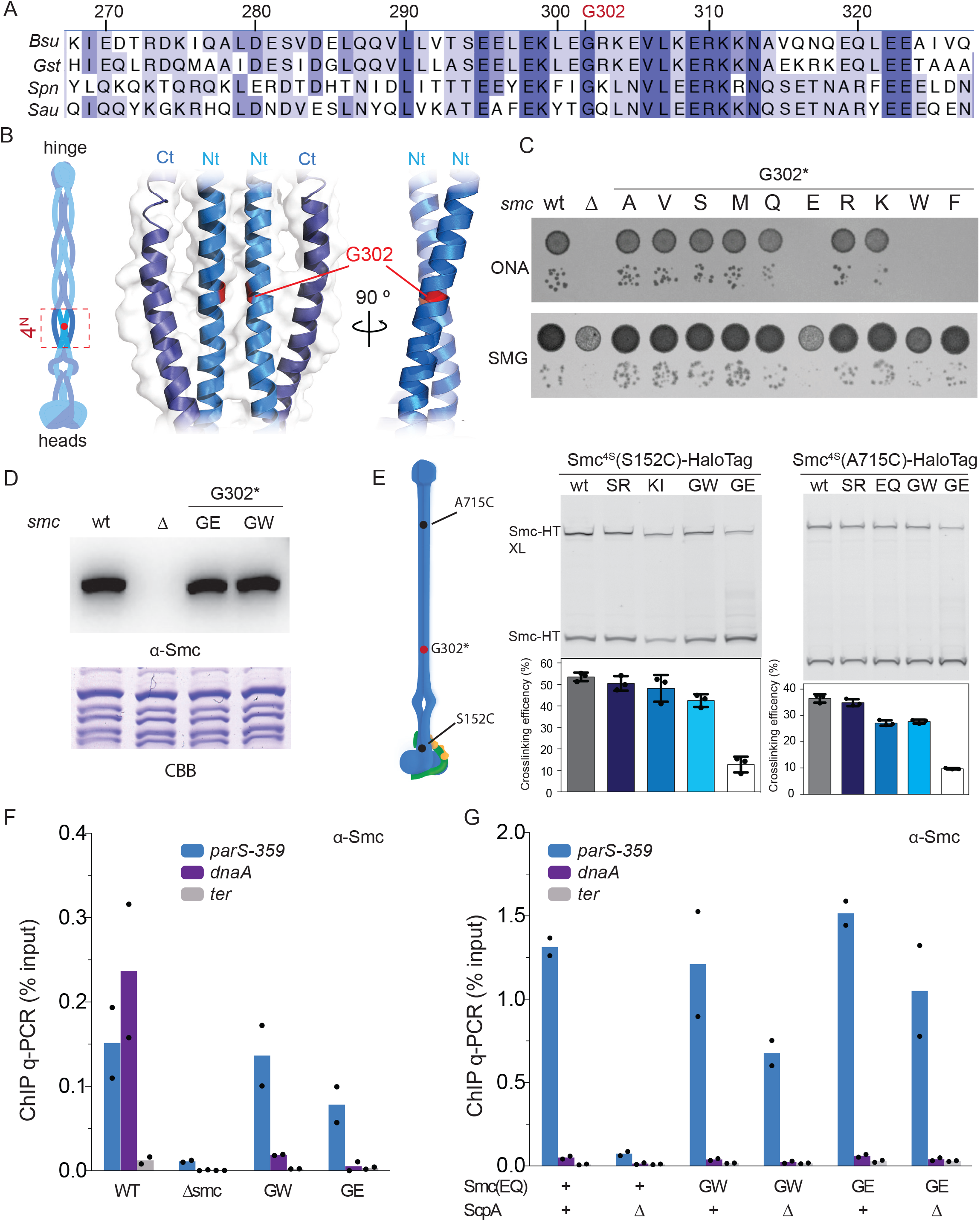
Residue G302 at the 4N arm contact is essential for Smc function. A. Alignment of 4^N^ arm contact sequences from selected firmicute species (*Bsu, Bacillus subtilis; Gst, Geobacillus stearothermophilus; Spn, Streptococcus pneumoniae; Sau, Staphylococcus aureus*). Residues G302 is indicated. B. The position of residue G302 at the 4^N^ arm contact (PDB: 5NNV). Left panel: schematic of arm contact zones as in Figure 2B; right panel: structural view of the 4^N^ arm contact from *Bsu* Smc in cartoon representation in front and side views. Colours as in Figure 3B. G302 is shown in red colours. C. Colony formation by *Bsu* strains with single-amino acid substitution in G302. 9^2^-fold and 9^5^-fold dilutions of stationary phase cultures were spotted on nutrient-rich medium (ONA) and nutrient-poor minimal glucose-glutamate medium (SMG) and grown at 37°C for 16 and 24 h, respectively. D. Smc protein levels in mutants G302E (GE) and G302W (GW) grown in SMG medium determined by immunoblotting with polyclonal antibodies raised against *Bsu* Smc protein. Coomassie staining (CBB) of cell extracts on separate gels shown as control for uniform protein extraction. E. Quantification of cross-linked species of Smc^4S^-HaloTag (C119S, C437S, C826S, C1114S) harbouring G302 mutations (labelled with TMR-HaloTag ligand). Cysteine pairs were introduced at reporter positions S152 (left panels) and A715 (right panels). Cross-linking efficiencies were calculated by in-gel fluorescence quantification of band intensities of non-crosslinked (Smc-HT) and crosslinked (Smc-HT XL) species. Efficiencies were compared between wild type (wt), the ABC signature motif mutant S1090R (SR), the ATP-binding mutant K37I (KI), G302W (GW), and G302E (GE) mutants. F. Chromatin-immunoprecipitation coupled to quantitative PCR (ChIP-qPCR) in *Bsu* cells using α-Smc antiserum. Selected loci near the replication origin and at the terminus regions were analysed. Dots represent data points from one out of two biological replicates. Mean values are indicated as bars. G. Chromatin-immunoprecipitation coupled to quantitative PCR (ChIP-qPCR) using α-Smc antiserum. As in (F) for strains carrying the E1118Q (EQ) mutation and lacking or having *scpA*.

### Defective chromosome loading of 4^N^ arm contact mutants

We next determined the chromosomal distribution of Smc(GE) and Smc(GW) proteins by chromatin-immunoprecipitation (ChIP) using α-ScpB and α-Smc antisera. Both mutants had close to normal enrichment at the *parS-359* site but very low enrichment at the *dnaA* locus as determined by ChIP-qPCR (Figure 4F). These mutants thus appear to target normally to *parS* but then fail to load onto DNA or to translocate along DNA (Minnen et al., 2016). Consistent with this notion, we found that their E1118Q variants showed elevated levels of enrichment at *parS*, similar or even higher than the otherwise wild-type Smc(E1118Q) protein. Formation of the 4^N^ arm contact may thus be required for the loading reaction or for the release of Smc protein from ParB/*parS* loading sites. Notably, a putative ParB/Smc interface has been mapped onto the Smc arm in the vicinity of the 3^C^ and 4^N^ arm contacts (Minnen et al., 2016).

We found that the GW and the GE mutation alleviated the requirement for ScpA in the targeting of the Smc(E1118Q) protein to a *parS* site (Figure 4G). ScpA may thus contribute to overcoming constraints imposed by the aligned Smc arms to facilitate the transition to the open conformation for *parS* targeting (Minnen et al., 2016). The G302 mutations likely alleviate these constraints and eliminate the requirement for ScpAB. The integrity of the 4^N^ arm contact is subsequently required for the closure of the S compartment and the clearance of Smc-ScpAB from ParB/*parS*. Purified preparations of Smc(G302W) and to a weaker extent also Smc(G302E) displayed increased ATP hydrolysis activity, which was curiously hindered rather than enhanced by the addition of ScpAB and DNA (Figure S4C). These point mutations in the Smc arms thus lead to mis-regulation of the ATPase by DNA.

### Intermediary conformations *in vivo*

We next focused on the organization of Smc heads *in vivo* by inferring their conformations from patterns of cross-linking obtained with Smc(K1151C). The K1151C residue was previously engineered to detect the ATP-engaged state (Cα-Cα distance: 11.6 Å) by BMOE cross-linking (Minnen et al., 2016). Here, we made use of bis-maleimide cross-linkers with longer spacers— which supported robust cross-linking in intact *Bsu* cells—to detect alternative conformations (Figure 5A). We observed that cross-linking of Smc(K1151C)-HaloTag was much more efficient with BM^3^ (17.8 Å) and BM^11^ (62.3 Å) than with BMOE (8 Å) or BM^2^ (14.7 Å) (Figure 5A and S5A) (Minnen et al., 2016). The Walker A ATP-binding mutant Smc(K37I) mutant also showed an upward trend in cross-linking with extending spacer length, but the cross-linking efficiencies did not reach levels comparable to wild-type Smc (Figure 5A). The Smc(E1118Q) protein displayed roughly equivalent cross-linking efficiencies regardless of spacer length (Figure 5A and S5A). Finally, a clear downward trend in cross-linking with increasing spacer length was observed when the E1118Q mutation was combined with three different Smc arm alterations (Figure 5B). For example, the Smc(204Ω/996Ω, E1118Q) protein—harboring a 13 amino acid peptide insertion in each arm—showed most efficient cross-linking with BMOE and decreasing efficiencies with increasing spacer length (Diebold-Durand et al., 2017). A similar trend was observed with an artificially shortened ‘mini’-Smc(E1118Q) protein (Bürmann et al., 2017) and with Smc(GE, E1118Q) (Figure 5B). This downward trend in cross-linking was expected for the ATP-engaged state because the K1151C residues are closely juxtaposed and thus ideally positioned for cross-linking by the smallest compound BMOE (Minnen et al., 2016).

**Figure 5.**
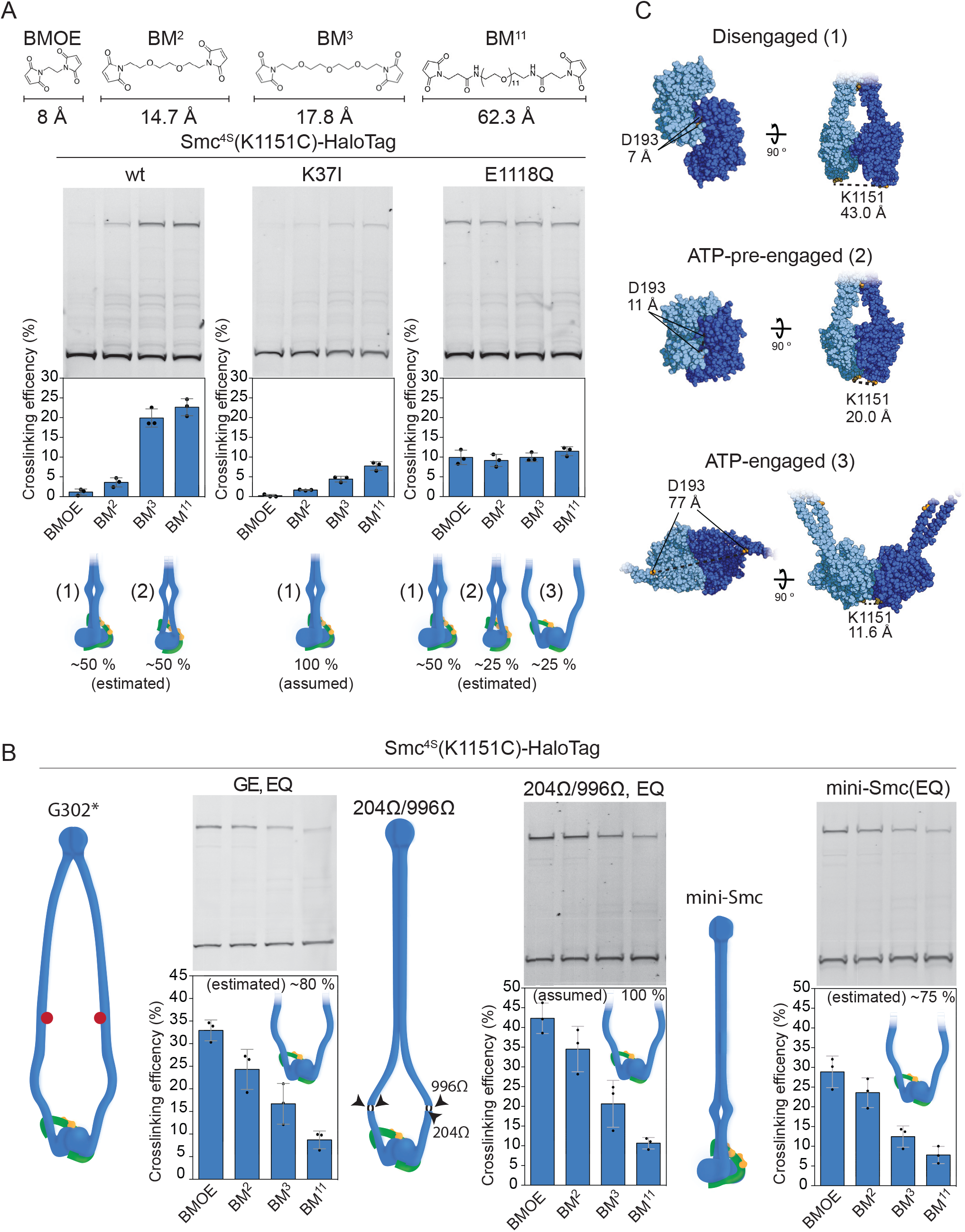
Smc head cross-linking *in vivo*. A. Top panel: Chemical structures of bis-maleimido crosslinkers with variable spacer arms (BMOE, bismaleimidoethane; BM^2^, 1,8-bismaleimido-diethyleneglycol; BM^3^, 1,11-bismaleimido-triethyleneglycol; BM^11^, Bis-MAL-dPEG11). Bottom panels: Quantification of cross-linked species of Smc^4S^ harbouring the K1151C cysteine pair reporter using different crosslinkers. Crosslinking efficiencies were calculated as described for Figure 4E and compared between wt (left panel), K37I (middle panel) and E1118Q (right panel) variants. Error bars were calculated as standard deviation from 3 biological replicates. The values and schematics shown below the graphs indicate the approximately estimated relative abundance of different conformations, based on assumed abundances for the K37I mutant and the Smc(204Ω/996Ω, EQ) protein (panel B). B. Smc(K1151C) crosslinking in strains harbouring arm modifications. G302E (GE) (left panel), unstructured peptide insertions 204Ω/996Ω (middle panel) (Diebold-Durand et al., 2017) and mini-Smc CC298 (Bürmann et al., 2017) (right panel). Representation and quantification as in (A). C. Structural models of different ATPase states. Model for the disengaged state (1) as described in (Diebold-Durand et al., 2017). The ATP-engaged state (3) is based on a crystal structure (PDB: 5XG3). The ATP-pre-engaged state (2) is manually built to aligning ABC motifs without opening the arms. Individual subunits are coloured in light or dark blue colours, respectively. D193 and K1151 inter-subunit Cα-Cα distances are shown as a reference for the different conformations.

We conclude that in addition to two previously known states (disengaged and ATP-engaged) (Figure 1A), Smc heads also occur in a third, hitherto uncharacterized state. We envision that in this ‘ATP-pre-engaged’ state active site residues from both heads are aligned (thus supporting efficient K1151C cross-linking by BM^3^ and BM^11^), but the closure of the catalytic pocket remains incomplete due to a gap persisting between the signature motif residues from one head and the ribonucleotide bound to the opposing head (thus reducing K1151C cross-linking by BMOE). Taking all *in vivo* cross-linking data into account (Figure 5A and B), we next estimated the occupancy of the different states in wild-type Smc. Assuming a 100 % occupancy of the ATP-engaged state in Smc(204Ω/996Ω, E1118Q) (Figure 5B) and of the disengaged state in the ATP-binding mutant Smc(K37I) (Figure 5A), we propose that wild-type Smc-ScpAB complexes rarely populated the ATP-engaged state and distributed roughly equally between the other states (Figure 5A). In Smc(E1118Q), about a quarter of complexes appear to be ATP-engaged suggesting that head engagement is an inefficient and stepwise process *in vivo*, unless arm integrity is affected.

Of note, while ScpA is crucial for forming the ATP-engaged state in Smc(E1118Q), the requirement for ScpA was (partially) alleviated in the GW and GE variants (Figure S5E) (Minnen et al., 2016). As mentioned above, ScpA was dispensable for the targeting of the GE and GW Smc(E1118Q) proteins to *parS* sites (Figure 4G) being consistent with the notion that ScpAB helps to disrupt arm co-alignment to facilitate head engagement and *parS* targeting. This is in excellent agreement with the efficient and ScpA-independent *parS* targeting of a hinge dimerization mutant of Smc(EQ), which also displays mis-aligned Smc arms and increased levels of head engagement (Minnen et al., 2016).

Finally, we wanted to establish whether the ATP-pre-engaged state—detected by BM^3^ and BM^11^ cross-linking of Smc(K1151C)—corresponded to the closed, the open or the intermediary conformation. Using a combination of cysteine residues, we found that arm cross-linking with Q320C (at contact 4^N^) appeared to occur largely independently of K1151C cross-linking, implying that the 4^N^ arm contact is mostly engaged when heads are cross-linked at K1151C (Figure 6A). The efficient BM^3^ cross-linking of ATP-pre-engaged heads (in otherwise wild-type Smc) thus must have occurred on a (largely) closed conformation, implying that the closed conformation exists in a disengaged and an ATP-pre-engaged state (Figure 6B).

**Figure 6.**
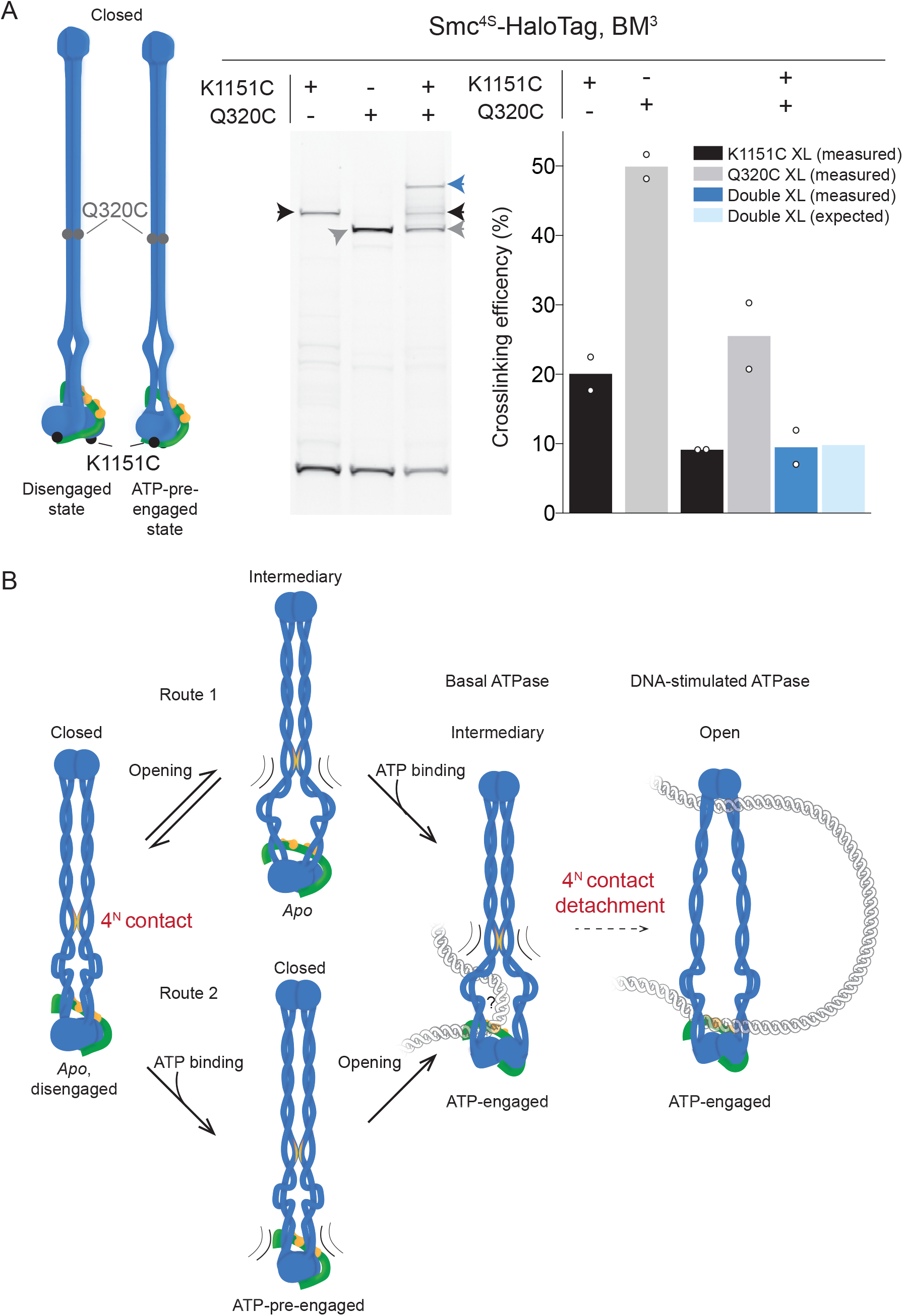
A. Simultaneous cross-linking of *Bsu* Smc heads and arms *in vivo*. Left panel: Schematics of Smc-ScpAB in closed conformations with disengaged or ATP-pre-engaged heads. Cysteine reporters Q320C and K1151C are indicated as dots in black and gray colours, respectively. Right panel: Quantification of single and double cross-linking efficiencies in Smc^4S^-HaloTag with Q320C and/or K1151C. The grey arrowhead denotes the species cross-linked only at Q320C; species only cross-linked at K1151C (black colours); species cross-linked at Q320C and K1151C (blue colours). Individual data points are shown as white circles from two biological replicates. Average values are shown as bars (same colour coding as for arrowheads). Data is also shown in Figure S6. B. Model of the conformational landscapes and multi-step opening of Smc-ScpAB. Two alternative pathways in the basal ATPase. Route 1: Spontaneous separation of the juxtaposed heads leads to an intermediary conformation with constraint arms, which is then stabilized by ATP binding and head engagement. Route 2: ATP binding in the closed conformation generates ATP-pre-engaged heads, which are constraint by the arms. Partial arm opening and head engagement result in the intermediary conformation. Optional: Full opening of the arms produces the DNA-stimulated ATPase.

## Discussion

Chromosomes undergo continuous reorganization by DNA loop extrusion. The latter involves a unique DNA motor which moves by apparently making large translocation steps. Elucidating the underlying mechanism requires an understanding of the conformational transitions in SMC complexes during the ATP hydrolysis cycle as well as the concomitant DNA binding and unbinding events. Here we provide structural insights into three separate conformations and corresponding state of the ATPase. The conformational dynamics appears largely governed by a limited rigidity of the head-proximal Smc arms and by a finite stability of arm contacts. While the architecture appears dynamic, principles of organization can be deduced.

### The conformational landscape of SMC complexes

Cryo-electron microscopy recently helped to reveal the molecular architecture of SMC-kleisin complexes (in apo and ATP-engaged states) from eukaryotes (Collier et al., 2020; Higashi et al., 2020; Lee et al., 2020; Shi et al., 2020). Solving structures of more dynamic or flexible conformations, however, remains challenging even for cryo-EM. EPR is ideally suited to help close these gaps by providing insights into the distributions of conformations (Jeschke, 2018; Kazmier et al., 2014; Vercellino et al., 2017).

### The closed conformation

Independent lines of evidence point to the prevalence and conservation of a closed conformation in SMC complexes. It is predominant in reconstituted Smc-ScpAB, as judged by EPR (Figure 1), and in living *Bsu* cells according to site-specific cross-linking experiments (Diebold-Durand et al., 2017). Its main characteristic features are disengaged heads, and completely aligned arms. Point mutations that hinder arm alignment eliminate Smc function in *Bsu* implying that the closed conformation is critically important (Figure 4). Yeast condensin was found by cryo-electron microscopy as an elongated particle with ‘non-engaged’ heads and completely aligned arms (Lee et al., 2020), presumably corresponding to the Smc-ScpAB rod, albeit displaying folded Smc2 and Smc4 arms (see below). Intriguingly, the Smc2 and Smc4 arms in *apo* condensin are most closely juxtaposed at the region that corresponds to the 4^N^ arm contact in Smc-ScpAB, being consistent with the idea that this contact has retained its special function in yeast condensin, and possibly other SMC complexes, too (Figure S4D). The condensin arms bend ∼180° at an elbow allowing the hinge to fold back and contact the middle of the Smc2 arm (Lee et al., 2020). This feature has not been observed for Smc-ScpAB and may have evolved in eukaryotic SMC proteins (and independently also in MukBEF) (Bürmann et al., 2019). The function of arm folding and of the hinge/arm contact is not well understood. It is conceivable that the folding may be relevant to the process of stepwise opening and closing of the S compartment, possibly stabilizing half-open states and thus providing additional levels of control. Of note, the closed conformation is rare in AFM images of yeast condensin and *Bsu* Smc-ScpAB, unless the material is cross-linked prior to sample deposition (Fuentes-Perez et al., 2012; Ryu et al., 2019; Yoshimura et al., 2002). We suspect that the nonspecific adsorption of samples onto the AFM mica may affect SMC rod folding dynamics.

### The open conformation

The open conformation of Smc-ScpAB has previously been inferred indirectly from crystal structures of isolated ATP-engaged heads and supported by low resolution images obtained by rotary shadowing electron microscopy and by loss of cysteine cross-linking (Anderson et al., 2002; Diebold-Durand et al., 2017; Kamada et al., 2017; Lammens et al., 2004). In this study, we provide direct evidence for the existence of the open conformation. We characterize its shape and show that it represents an integral part of the DNA-stimulated ATP hydrolysis cycle. Smc dimers that are unable to completely open the S compartment retained the ability to hydrolyze ATP but lost the stimulation upon DNA addition. Hinge/DNA binding may open the arms at the hinge as previously proposed (Soh et al., 2015), thus facilitating the transition to ATP-engaged heads and supporting efficient ATP hydrolysis.

### Intermediary conformations

Here, we have expanded our knowledge on the conformational landscape of Smc-ScpAB by detecting and characterizing intermediary conformations using three complementary approaches (EPR, ATPase measurements and cysteine cross-linking) (Figure 1, 2 and 5). Altogether, our data suggest that Smc heads and the head-proximal arms undergo dynamic structural changes.

Patterns of cross-linking demonstrate that the Smc heads can adopt at least three different states *in vivo*, here denoted as disengaged, ATP-pre-engaged, and ATP-engaged state (Figure 6B). The ATP-engaged state occurs in the open and the intermediary conformation. We propose that in the ATP-pre-engaged state, the active site residues from both heads are aligned via ATP but the arms remain (largely or completely) aligned. The heads may transition gradually from the disengaged, via the ATP-pre-engaged to the ATP-engaged state. Both steps might be impeded by arm alignment.

Artificially engineered arm junctions reduce—but do not block—ATP hydrolysis (Figure 2 and 3). The ATP-engaged state—presumably a prerequisite for ATP hydrolysis—is thus compatible with partially open arms. Whether such basal ATPase cycles are futile or contributing to DNA translocation remains to be established. The fact that the essential head/DNA interfaces is exposed in the intermediary conformation (while the non-essential hinge/DNA interface is likely inaccessible) is consistent with the notion that this state is functionally relevant (Figure 6B), (Vazquez Nunez et al., 2019).

Cryo-EM structures of DNA-bound cohesin in the ATP-engaged state show a partially open conformation with separated head-proximal arms (Collier et al., 2020; Higashi et al., 2020; Shi et al., 2020). Most of the more distal arm sequences are less well resolved—probably due to structural flexibility—but appear to remain aligned (and elbow-folded) (Collier et al., 2020; Higashi et al., 2020). Whether conformations with completely dissociated arms are present in cohesin (and condensin) is not fully established, but rotary-shadowing electron micrographs would be consistent with this notion (Anderson et al., 2002). Upon arm dissociation, the arms either remain folded at the elbow thus producing B-shaped structures as inferred from AFM images of condensin (Eeftens et al., 2016) or they unfold at the elbow to generate O-shaped complexes (as seen by EM and similar to the open conformation of Smc-ScpAB).

The intermediary conformations of Smc-ScpAB are structurally not well defined, probably due to significant flexibility in the head-proximal arms. In condensin, however, a well-defined partially open state was detected by cryo-EM. This *apo* bridged conformation of yeast condensin showed partial separation of arms and heads (Lee et al., 2020). The separated heads are however bridged—and likely stabilized—by one of two hawk subunits, which are specific to condensin and cohesin. Our observations, together with the recent cryo-EM structures, indicate that intermediary conformations (with partially opened arms) are widely conserved in SMC complexes.

### The DNA translocation cycle

It is tempting to speculate that the intermediary conformations control the process of compartment opening and closure and direct the binding and unbinding of suitable DNA substrates thus driving directional DNA translocation. Here, we propose that one of the eight arm contacts acts as a molecular switch for arm opening in response to initial DNA contact (Figure 6B). Chromosomal DNA is entrapped in the K compartment of the closed conformation as determined by site-specific crosslinking in *Bsu* Smc-ScpAB (Vazquez Nunez et al., 2019) and in yeast cohesin (Chapard et al., 2019). The open conformation holds two DNA binding interfaces in the S compartment, of which the head/DNA interface—but not the hinge/DNA interface—is essential for Smc function and critical for Smc DNA translocation (Hirano and Hirano, 2006; Vazquez Nunez et al., 2019). These observations suggest that DNA is transiently occupying the S compartment. How DNA enters and exits the S compartment is a crucial open question.

The loop-capture model proposes that DNA is inserted as a loop on top of the heads and upon ATP hydrolysis exits the S compartment by transfer to the K compartment between the heads (Diebold-Durand et al., 2017; Marko et al., 2019). Alternatively, the DNA enters the S compartment by transfer between the heads as recently proposed for yeast cohesin (Collier et al., 2020). How the latter may lead to DNA translocation and DNA loop extrusion is however unclear. Unilateral and gradual opening of the arms—as described in this study—is likely directly relevant to this key substrate selection process. Future experiments will have to establish how the conformations (open, closed and intermediary) of SMC complexes associate with chromosomal DNA double helices to promote DNA loop extrusion.

## Acknowledgements

We thank Frank Bürmann for help in setting up initial EPR-DEER experiments and for helpful comments on the manuscript and Daniel Klose for collecting EPR-DEER data. We are grateful to all members of the Gruber lab for support and stimulating discussions. We thank the Genome Technologies Facility (GTF) at the University of Lausanne for deep sequencing. This work was supported by the European Research Council (Horizon 2020 ERC CoG 724482).

## Methods

### Experimental Model

#### *Bacillus subtilis* strains and growth

*Bsu* strains used in this work are derived from the 1A700 isolate. Genotypes and strain numbers are listed in Table S5. Strain usage is detailed in Table S1. Naturally competent *Bsu* cells were transformed as in (Diebold-Durand et al., 2017) including a longer starvation incubation time (2 hours) for high efficiency. The transformants were selected on SMG-agar plates with appropriate antibiotics and single-colonies were isolated. The strains were confirmed by PCR and Sanger sequencing as required. For dilution spot assays, cells were grown for 8 hours at 37 °C in SMG medium and then were diluted to 81 and 59 000-fold. Afterwards, dilutions were spotted onto ONA (16 h incubation) or SMG (24 h incubation) agar plates at 37 °C (Vazquez Nunez et al., 2019).

### Protein purification

#### Full-length *Bsu* Smc

Native Smc proteins were purified following the procedure described in (Bürmann et al., 2017). *E. coli* BL21-Gold (DE3) strain was transformed with pET-22 or pET-28 derived plasmids containing the *smc* recombinant sequences. Proteins were expressed using ZYM-5052 autoinduction medium for 23 h at 24 °C. Cells were resuspended in lysis buffer (50 mM Tris-HCl pH 7.5, 150 mM NaCl, 1 mM EDTA, 1 mM DTT, 10 % (w/v) sucrose) supplemented with protease inhibitor cocktail (PIC) and sonicated. The lysate was centrifuged, the supernatant was filtered with a 0.45 µM pore size membrane and then loaded onto two HiTrap Blue HP 5 mL columns connected in series and eluted with lysis buffer containing 1 M NaCl. The main peak elution fractions where diluted in salt-less buffer (50 mM Tris-HCl pH 7.5, 1 mM EDTA, 1 mM DTT) to a conductivity equivalent of 50 mM NaCl (≈ 8 mS/cm). The diluted sample was supplemented with PIC and loaded on a HiTrap Heparin HP 5 mL column. Elution was performed by applying a linear gradient of buffer up to 2 M NaCl. The main peak fractions (aprox. 5 mL) were collected and further purified by gel filtration on a XK 16/70 Superose 6 PG column in 50 mM Tris-HCl pH 7.5, 200 mM NaCl, 1 mM EDTA, 1 mM TCEP. Main peak fractions where collected, concentrated with a Vivaspin 15 10K MWCO filter, flash frozen with liquid nitrogen and stored at −80 °C. Protein concentration was calculated by absorbance using theoretical molar absorption and molecular weight values.

#### ScpA

Native ScpA was purified following the procedure reported in (Vazquez Nunez et al., 2019). The protein was expressed using *E. coli* BL21-Gold (DE3) transformed with a pET-28 derived plasmid containing ScpA coding sequence. Cells were cultivated in ZYM-5052 autoinduction medium at 16 °C for 28 h, harvested and resuspended in lysis buffer (50 mM Tris-HCl pH 7.5, 200 mM NaCl, 5 % glycerol) supplemented with PIC. After sonication and centrifugation, the supernatant was applied to a 5 mL HiTrapQ ion exchange column and eluted with a gradient up to 2 M NaCl. The peak fractions were mixed with 4 M NaCl buffer to reach a final concentration of 3 M NaCl. The sample was injected into a HiTrap Butyl HP column and eluted in a reverse gradient to 50 mM NaCl. Peak fractions were pooled and concentrated to 5 mL in Vivaspin 15 10K MWCO filters and subsequently purified by gel filtration in a Hi Load 16/600 Superdex 75 pg column, equilibrated in 20 mM Tris-HCl pH 7.5, 200 mM NaCl. Protein was concentrated, aliquoted, flash frozen and stored at −80 °C.

#### ScpB

Native ScpB was purified using the procedure described in (Vazquez Nunez et al., 2019). A pET-22 derived plasmid with the coding sequence of ScpB was transformed in chemically competent *E. coli* cells. Cells were cultivated I ZYM-5052 autoinduction medium at 24 °C for 23 h. Cells were harvested and resuspended in lysis buffer (50 mM Tris-HCl pH 7.5, 150 mM NaCl, 1 mM EDTA, 1 mM DTT) supplemented with PIC. After sonication and centrifugation, the supernatant was diluted to 50 mM NaCl, loaded onto a 5 mL HiTrap Q HP column and eluted with a gradient to 2 M NaCl. The sample was diluted in lysis buffer with 4 M NaCl buffer in order to reach a final concentration of 3 M NaCl. The sample was applied on two 5 mL HiTrap Butyl column connected in series. Protein was eluted with a reverse gradient to 50 mM NaCl. A second run on the 5 mL HiTrap Q HP column in the same conditions as described above was necessary in some cases when the sample still showed 260/280 ratios close to 1. The peak fractions were concentrated and further purified by SEC using a Hi Load 16/600 Superdex 200 pg equilibrated in 50 mM Tris-HCl pH 7.5, 100 mM NaCl, 1 mM DTT. The fractions containing the protein were concentrated, aliquoted, flash frozen and stored at −80°C.

### Method details

#### ATPase assay

ATPase activity measurements were done using the coupled reaction of pyruvate kinase/lactate dehydrogenase (Bürmann et al., 2017). ADP production was monitored for 1 h by oxidation of NADH absorbance changes at 340 nm. The values were collected using a Synergy Neo Hybrid Multi-Mode Microplate reader. The reaction mixture consisted in 1 mM NADH, 3 mM Phosphoenol pyruvic acid, 100 U Pyruvate kinase, 20 U Lactate dehydrogenase and the appropriate ATP concentrations. Double-stranded oligonucleotides (40 bp (5’-TTAGTTGTTC GTAGTGCTCG TCTGGCTCTG GATTACCCGC)) were added for measurements that required it to a final concentration of 3 µM, unless indicated differently. The final protein concentration in the assay was 0.15 μM Smc dimers in ATPase assay buffer (50 mM HEPES-KOH pH 7.5, 50 mM NaCl, 2 mM MgCl2). Measurements were carried out at 25 °C.

#### M3M crosslinking-ATPase assay

For this assay, a fresh full-length Smc purification (as described above) was done for each experiment. The last SEC purification step, however, was done in 50 mM Tris-HCl pH 7.5, 200 mM NaCl in order to remove any trace of reducing agents. The peak fractions were concentrated to ∼35 µM Smc dimer in a Vivaspin 15 10K MWCO filters. The crosslinked reaction was carried on by mixing 10-fold thiol molar excess of M3M (diluted in DMSO). A typical reaction was done in 500 µL with 30 µM Smc dimers and 600 µM M3M in 50 mM Tris-HCl pH 7.5, 200 mM NaCl at 4 °C overnight, protected from light. The final concentration sometimes differed slightly, depending on the purification yield of the single cysteine mutant, but the M3M:thiol ratio was always maintained. A negative control was always included by mixing the same amount of protein with an equivalent volume of DMSO as in the experimental M3M reaction.

After crosslinking the sample was centrifuged to remove big aggregates and the excess of crosslinker was removed with a Zeba spin 7K MWCO 0.5 mL desalting column, preequilibrated with 50 mM Tris-HCl pH 7.5, 200 mM NaCl. An additional gel filtration purification was done to remove small aggregates and traces of crosslinker in a Superose 6 increase 10/300 column equilibrated in 50 mM Tris-HCl pH 7.5, 200 mM NaCl. Afterwards, the fractions at the maximum of the peak were recovered. The concentration of the protein was typically ∼4 µM Smc dimer. The sample was mixed with ScpA and ScpB and the ATPase assay was performed as described above. 10 mM DTT final concentration was added for the samples that required it.

#### Fluorescence anisotropy measurements

Fluorescence anisotropy was measured using a 40 bp dsDNA (as described in ATPase assay) modified at the 3′ end with fluorescein. Measurements were recorded using a Synergy Neo Hybrid Multi-Mode Microplate reader (BioTek) with the appropriate filters in black 96-well flat bottom plates at 25°C. Buffer conditions were 50 mM Tris-HCl pH 7.5, 50 mM NaCl, 2 mM MgCl_2_ 50 nM dsDNA and 1 mM ATP for all measurements. Anisotropy measurements where exported from the BioTek Synergy Neo software and subsequently fit to a binding polynomial using non-linear regression in GraphPad Prism 8 (Vazquez Nunez et al., 2019)

#### Double Electron-Electron Resonance (EPR-DEER)

Smc^3C^, ScpA and ScpB purifications were performed as described above, except for the last gel filtration step of Smc purification which was performed in absence of reducing agent. In order to avoid premature thiol oxidation, the sample was mixed with 10-fold molar excess of MTSL per thiol, immediately after the last purification step. The reaction volume was 500 µL and consisted in ∼40 µM Smc dimer and 800 µM MTSL in 50 mM Tris-HCl pH 7.5 at 4 °C and 200 mM NaCl. The reaction was incubated over-night at 4 °C protected from light. After the incubation, the sample was centrifuged to remove big aggregates and the MTSL excess was removed using a Zeba Spin desalting column, 7K MWCO, 0.5 mL. Afterwards the sample was further purified by gel filtration in a Superose 6 10/300 increase, equilibrated with 50 mM Tris-HCl pH 7.5 at 4 °C, 200 mM NaCl and 90 % deuterium oxide (heavy water). The sample was concentrated to ∼20 µM Smc dimer and the label efficiency was determined by calculating the double integral of the continuous wave (CW) EPR spectrum. Room temperature CW EPR measurements were done at X band (∼ 9.7 GHz) using a commercial X-band Magnettech MiniScope MS 400. Spectra were taken at 13 dB attenuation, corresponding to 5 mW incident microwave power and 0.15 mT magnetic field modulation amplitude. Samples were placed into 1 mm o.d. glass capillaries. Spin labelling efficiency for all mutants was estimated by comparing double integrals of EPR signals from the Smc mutants and the concentration standard (200 µM water solution of TEMPOL).

Samples were flash frozen and stored at −80 °C until use. Before measurement, the protein samples (Smc, ScpA and ScpB) were thawed and centrifuged to remove big aggregates. The final concentration per condition were: 5 µM SmcScpAB, 10 µM dsDNA 40 bp and 1 mM ATP in 50 mM Tris-HCl pH 7.5 at 4 °C, 50 mM NaCl, 2 mM MgCl_2_, 40 % Ethylene glycol-d6 (as cryoprotectant) and 90 % deuterium oxide. Samples were incubated 5 minutes in 1.5 mL centrifuge tubes at 25 °C with gentile (700 rpm) shaking. Distance measurements were performed at Q-band frequencies (∼ 34.4 GHz) using a standard double electron-electron resonance (DEER) sequence (Pannier et al., 2000). DEER traces were acquired at 50 K. A commercial X/Q-band Elexsys E580 spectrometer (Bruker) power-upgraded to 200 W equipped with the homebuilt TE102 rectangular resonator was used (Polyhach et al., 2012). The protein solution ready for measurement was filled into the 3 mm o.d. quartz tubes, shock-frozen by immersion into liquid nitrogen and inserted into the pre-cooled resonator. All observer pulses were 16 ns long, the pump pulse was 12 ns long, the pump frequency was set at the global maximum of the nitroxide EPR spectrum and the observer frequency was set 100 MHz lower.

#### In vivo cross-linking

*Bsu* cultures in 200 mL SMG were grown to exponential phase (OD_600_ = 0.02) at 37 °C. Cells were collected by filtration harvesting and washed in cold PBS + 0.1 % (v/v) glycerol (‘PBSG’). A biomass equivalent to 0.85 OD units was sorted. Cells samples were centrifuged 2 min at 10,000 *g*, resuspended in fresh PBSG (200 µL). The cross-linking reaction was started by adding 0.5 mM BMOE and it was incubated for 10 min on ice. The reaction was quenched by the addition of 1.4 mM 2-mercaptoethanol. Cells were centrifuged and resuspended in 30 μL of lysis mix (75 U/mL ReadyLyse Lysozyme, 750 U/mL Sm DNase, 5 μM HaloTag TMR Substrate and protease inhibitor cocktail ‘PIC’ in PBSG). Lysis reaction was incubated at 37 °C for 30 min. Afterwards, the material was mixed with 10 μL of 4X LDS-PAGE buffer, samples were incubated for 5 min at 95 °C and resolved by SDS-PAGE. Gels were imaged on an Amersham Typhoon scanner with Cy3 DIGE filter setup.

#### Chromatin Immunoprecipitation

The procedure followed the one described in (Vazquez Nunez et al., 2019). *Bsu* strains were cultured in 200 mL SMG medium from OD600 = 0.004 initial OD to OD600 = 0.02 at 37 °C. Cells were crosslinked with 20 mL of buffer F (50 mM Tris-HCl pH 7.4, 100 mM NaCl, 0.5 mM EGTA pH 8.0, 1 mM EDTA pH 8.0, 10 % (w/v) formaldehyde) and incubated for 30 min at room temperature with occasional manual shaking. Cells were harvested by filtration and washed in PBS at 4 °C. The cell biomass equivalent to 2 OD_600_ units was resuspended in 1 mL TSEMS (50 mM Tris pH 7.4, 50 mM NaCl, 10 mM EDTA pH 8.0, 0.5 M sucrose and protease inhibitor cocktail) containing 6 mg/mL lysozyme from chicken egg white. Primary lysis was done at 37 °C for 30 min with shaking at 1400 rpm. Protoplasts were washed twice and resuspended in 1 mL TSEMS. The sample then was split into 3 aliquots and pelleted. Pellets were flash frozen and stored at −80 °C until used.

Each pellet was thawed and resuspended in 2 mL buffer L (50 mM HEPES-KOH pH 7.5, 140 mM NaCl, 1 mM EDTA pH 8.0, 1 % (v/v) Triton X-100, 0.1 % (w/v) Na-deoxycholate) containing 0.1 mg/mL RNase A and PIC. The suspension was sonicated during 1 min in 3 cycles of 20 seconds and 40 % amplitude at 4 °C using a MS73 probe. The sonicated material was centrifuged at 4 °C and 20,000 × g and 200 μL of the supernatant were collected as input reference. The immunoprecipitation was carried out by mixing 750 μL of the extract with 50 μL of Dynabeads Protein-G suspension freshly charged with 50 μL α-ScpB antiserum and incubated for 2 h on a wheel at 4 °C. Beads were washed with 1 mL each of buffer L, buffer L5 (buffer L containing 500 mM NaCl), buffer W (10 mM Tris-HCl pH 8.0, 250 mM LiCl, 0.5 % (v/v) NP-40, 0.5 % (w/v) Na-Deoxycholate, 1 mM EDTA pH 8.0) and buffer TE (10 mM Tris-HCl pH 8.0, 1 mM EDTA pH 8.0). All wash steps were done at 25 °C during 5 min shaking (1400 rpm). Beads were resuspended in 520 μL buffer TES (50 mM Tris-HCl pH 8.0, 10 mM EDTA pH 8.0, 1 % (w/v) SDS). The reference sample was mixed with, 300 μL buffer TES and 20 μL 10 % SDS. Formaldehyde cross-links were reversed over-night at 65 °C with vigorous shaking.

For phenol/chloroform DNA extraction, samples were cooled to room temperature, mixed with 500 μL phenol equilibrated with buffer (10 mM Tris-HCl pH 8.0, 1 mM EDTA), emulsified and centrifuged for 10 min at 20,000 × g. Subsequently, 450 μL of the supernatant was emulsified with 450 μL chloroform and centrifuged for 10 min at 20,000 × g. DNA precipitation was done by taking 400 μL of the supernatant and mixing with 1.2 μL GlycoBlue, 40 μL of 3 M Na-Acetate pH 5.2 and 1 mL ethanol and incubated for 20 min at −20 °C. Samples were centrifuged at room temperature and 20,000 × g for 10 min, and the precipitate was dissolved in 100 μL buffer PB (QIAGEN) for 10 min at 55 °C, purified with a PCR purification kit, and eluted in 50 μL buffer EB.

For qPCR, samples were diluted in DNase-free water (1:10 for IP and 1:1000 for input) and 10 μL reactions (5 μL Takyon SYBR MasterMix, 1 μL of 3 μM primer mix, 4 μL sample) were run in duplicates in a Rotor-Gene Q machine (QIAGEN) using the appropriate primer pairs.

### Quantification and statistical analysis

#### Analysis of cross-linking efficiencies

TMR fluorescence or Coomassie stained bands were quantified using ImageQuant TL 1D V8.1. Lanes were defined manually. Bands were detected automatically, and the band intensities corrected for background signal using the Rolling Ball algorithm with a ball radius set to 129. Values from three replicate experiments were exported to Microsoft Excel for calculation of average fractions and standard deviation.

#### Steady-state kinetics

The absorbance values were exported and fit to a straight-line equation in GraphPad prism 8. The slope values were transformed to rate values using the molar absorption coefficient of NADH. The rates were expressed into absolute values by correcting for the protein concentration. When an ATP titration was done, the rates were fit using non-linear regression to the Hill equation:

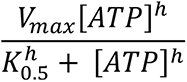

Where *V_max_* is the maximal rate at a given protein concentration, [ATP] is the variable ATP concentration, *K_0.5_* is the semi-saturation concentration and *h* is the Hill coefficient. Fit parameters obtained from the average of duplicates summarized in Table S4.

#### DEER traces analysis

Experimental DEER traces were processed with the open-source DeerAnalysis package (Jeschke et al., 2006), available at www.epr.ethz.ch/software/index. For background correction, a homogeneous distribution of spins in 3D space was assumed in all cases. Distance distributions for all mutants were calculated using double Gaussian model, whereas preliminary (calibrating) analysis for one of the mutants (D193) was performed model-free with the Tikhonov regularization.

### Analysis of qPCR Data

The threshold cycle (TC) was obtained by analyzing the fluorescence raw data in the Real-Time PCR Miner server (http://ewindup.info/miner/) (Zhao and Fernald, 2005). IP/input ratios were calculated as α 2ΔCT, where ΔCT = CT (Input) – CT and α is a constant determined by extraction volumes and sample dilutions. Data are presented as the mean of duplicates.

## Supplemental data

**Figure S1.**
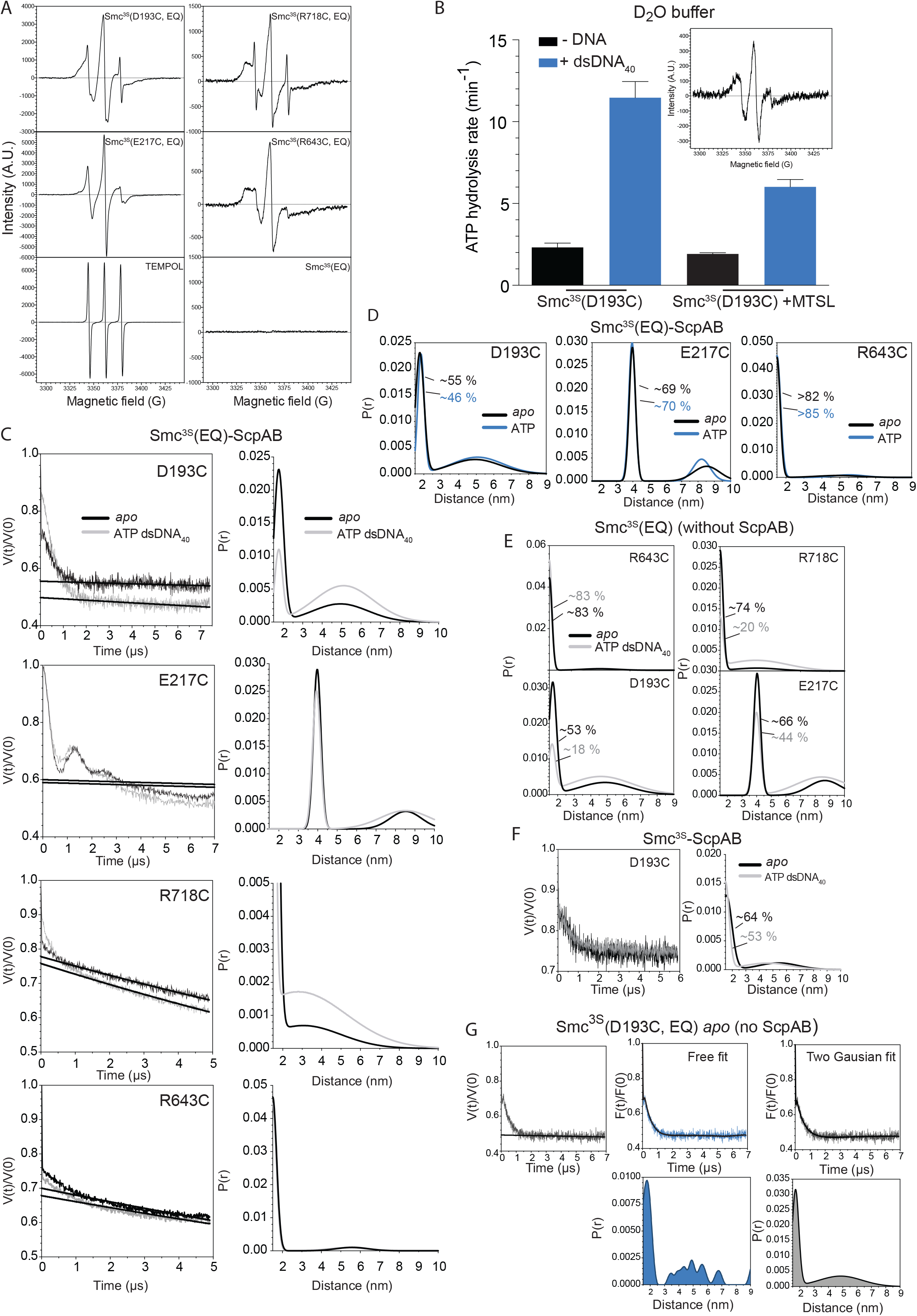
EPR-DEER analysis of Smc^3S^ variants. A. Continuous wave (CW) EPR spectra for Smc^3S^(EQ) single cysteine mutant proteins. Spin labelling efficiency for all mutants was estimated by comparing double integrals of EPR signals with the concentration standard (200 µM water solution of TEMPOL). Traces were obtained directly from the main peak elution fraction after MTSL excess removal (see methods). B. ATP hydrolysis rate of Smc^3S^(D193C) at 150 nM protein concentration and stoichiometric amounts of ScpAB_2_, in the presence of 1 mM ATP and with or without 3 µM dsDNA_40_. Measurements were done in deuterated environment (D_2_O) and with 40 % dEthylene glycol in order to mimic the DEER measurement conditions. Inset shows the CW-EPR spectrum as in (A). C. Primary DEER traces obtained from Q band measurements (V(t)/V(0)) in the *apo* (black colours) and in the ATP-, DNA-bound state (gray colours). Straight black lines correspond to the estimated background decay. Right panels correspond the two-Gaussian fit probability distributions as shown in Figures 1B and 1D. D. Probability distributions of spin pair distances in single cysteine mutants of Smc^3S^(EQ) in the presence of ScpAB and the presence (blue colors) and absence (black colors) of 1 mM ATP. Shown as in Figure 1B and 1D. E. Probability distributions of spin pair distances in single cysteine mutants in the absence of ScpAB and the presence (blue colors) and absence (black colors) of 1 mM ATP and 3 µM dsDNA_40_. Shown as in Figure 1B and 1D. F. Primary DEER traces (left panel) and probability distributions (right panel) of spin pair distances in Smc^3S^(D193C) in the apo state (black colours) and ATP- and DNA-bound state (grey colours). Shown as in (C). G. Primary DEER data from Smc^3S^(D193C-EQ) *apo* in the absence of ScpAB. The primary trace (left panel) was corrected for background (F(t)/F(0)) and model-free fitted with the Tikhonov regularization to provide the distance distribution shown in blue colours in the lower panel. Additionally, a two-Gaussian fit provided the distance distribution shown as a gray area in the lower panel. Given the good general agreement and the difficult of Tikhonov regularization in dealing with very narrow and very broad features simultaneously, two-Gaussian fits were used for all other mutants and conditions.

**Figure S2.**
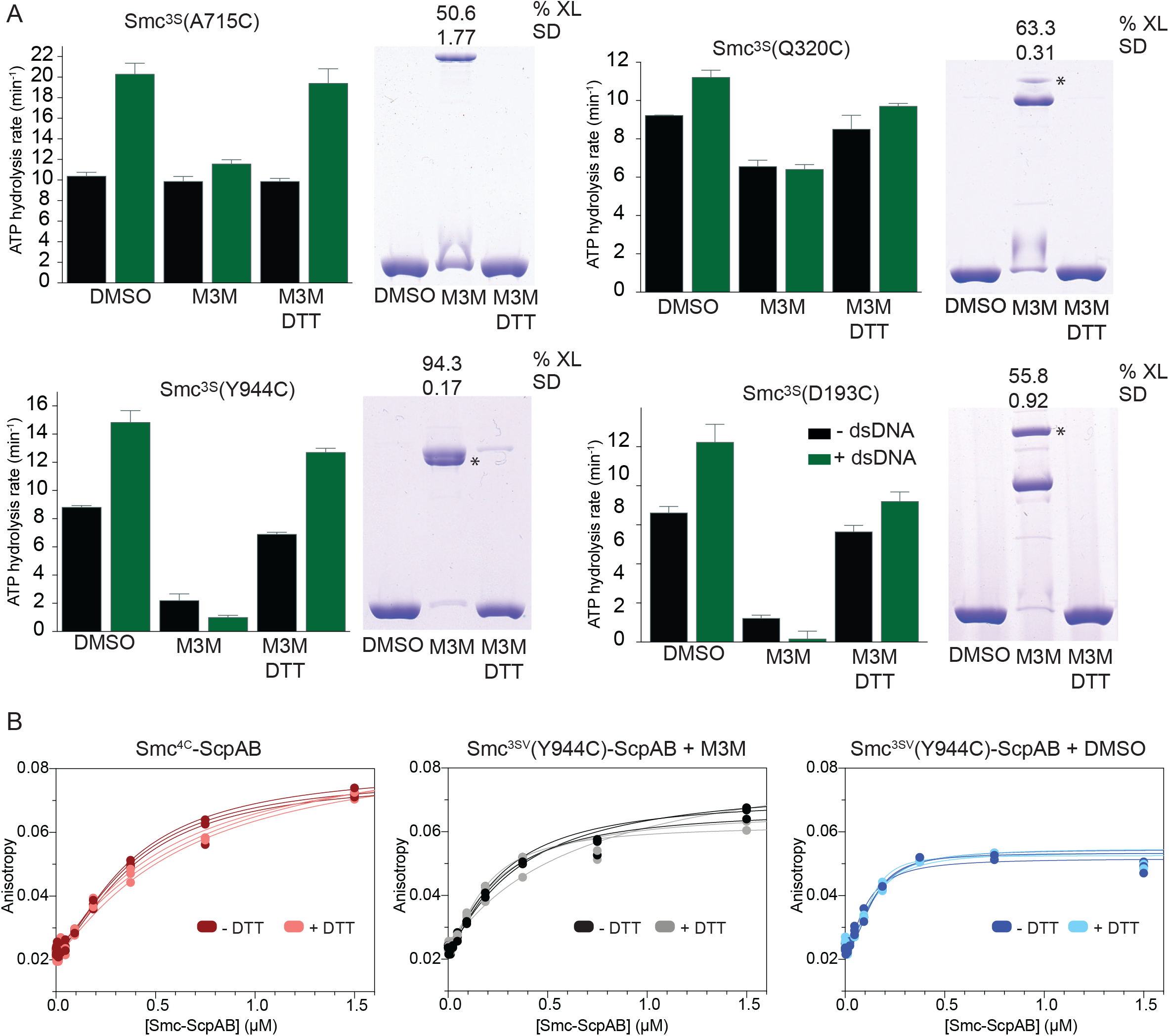
ATPase activity and DNA binding of M3M-cross-linked Smc^3S^ variants. A. Samples were prepared as described for Figure 2A. ATPase activity was determined with 1 mM ATP and measured in absence (black colours) and presence (green colours) of 3 µM dsDNA_40_. Error bars show the standard deviation from 3 technical replicates. Band intensities of CBB gel were quantified and M3M crosslink efficiency was determined. Mean and standard deviations are shown above each well calculated from 3 technical replicates. Bands derived from off-target cysteine cross-linking are labelled with asterisks. B. DNA binding of M3M-cross-linked Smc^3S^ variants as measured by fluorescence anisotropy with increasing protein concentration using fluorescein-labeled 40-bp dsDNA at 50 nM concentration. Data points from three experiments are shown as dots. The lines correspond to a nonlinear regression fit of the experimental data. Smc(Y944C) protein as in Figure 2C and also including wt Smc-ScpAB. Reduction of the M3M cross-link by addition of DTT had little effect on DNA binding.

**Figure S3.**
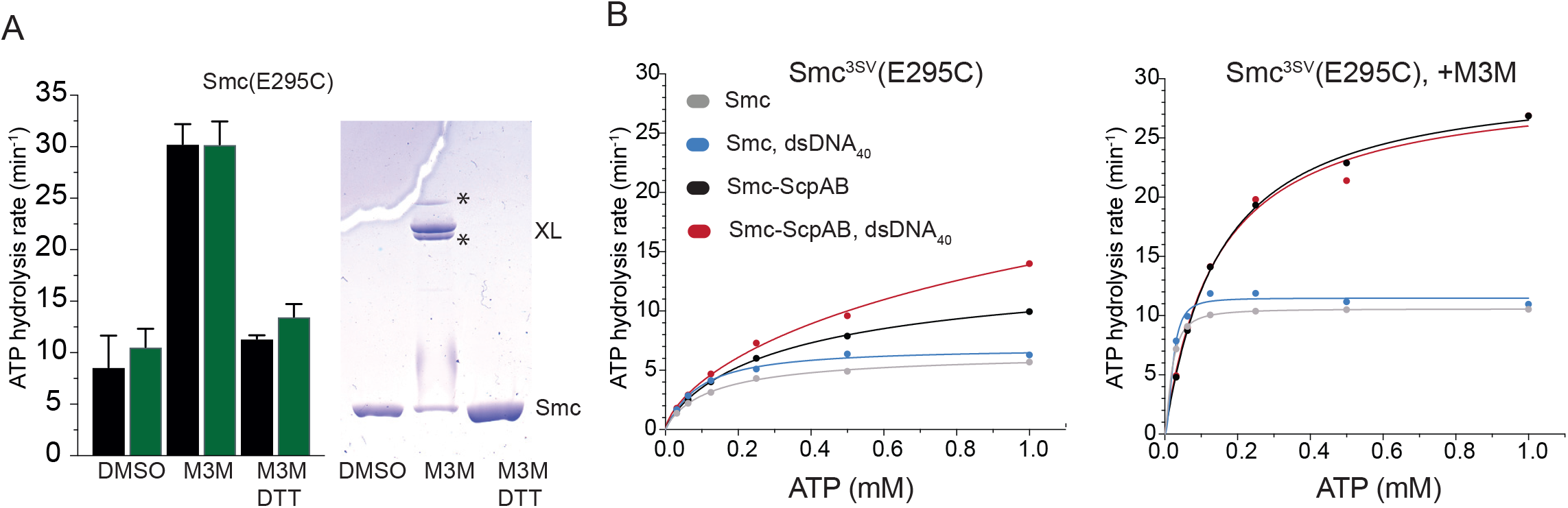
M3M crosslinking and ATPase activity of the Smc^3S^(E295C) variant. A. M3M crosslinking and ATPase activity of Smc^3S^(E295C) as in Figure S2A. B. ATP hydrolysis rates for Smc^3SV^(E295C) at variable ATP concentrations. As in Figure 2D.

**Figure S4.**
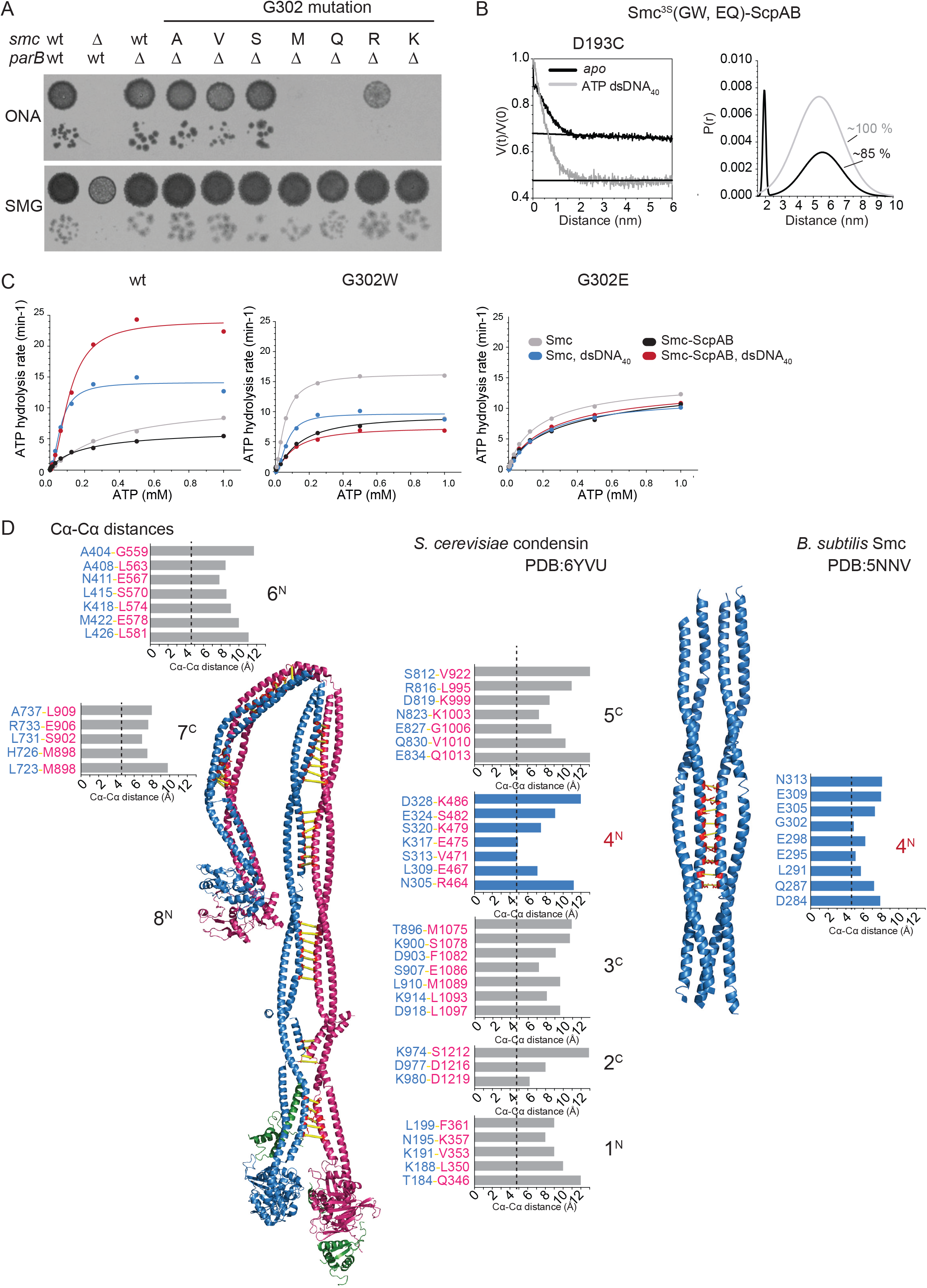
A. Dilution series spotting for *Bsu* strains carrying G302 single amino acid substitutions as well as a deletion of the *parB* gene. As in Figure 4C. B. Primary DEER traces (left panel) and probability distributions (right panel) of spin pair distances in Smc^3S^(D193C, G302W, E1118Q) in the apo (black colours) and ATP- and DNA-bound state. Shown as in Figure S1C. C. Measurement of ATP hydrolysis rates for Smc protein (wt), Smc(G302W) and Smc(G302E) with increasing concentrations of ATP and in the absence or presence of dsDNA_40_ and ScpAB. D. Cα-Cα distances measured at the inter-subunit interfaces. For *S. cerevisiae* condensin (PDB: 5YVU) (left panels). Smc2 (in blue colours), Smc4 (in pink colours) and Brn1 (in green colours) are shown in cartoon representation reported in the cryo-EM structure (Lee et al., 2020). Distances are displayed as bars in gray colours except for the 4^N^ arm contact, which are shown in blue colours. Dashed lines indicate the shortest distance observed as a reference. For the Bsu Smc arm crystal structure (PDB: 5NNV) (right panels).

**Figure S5.**
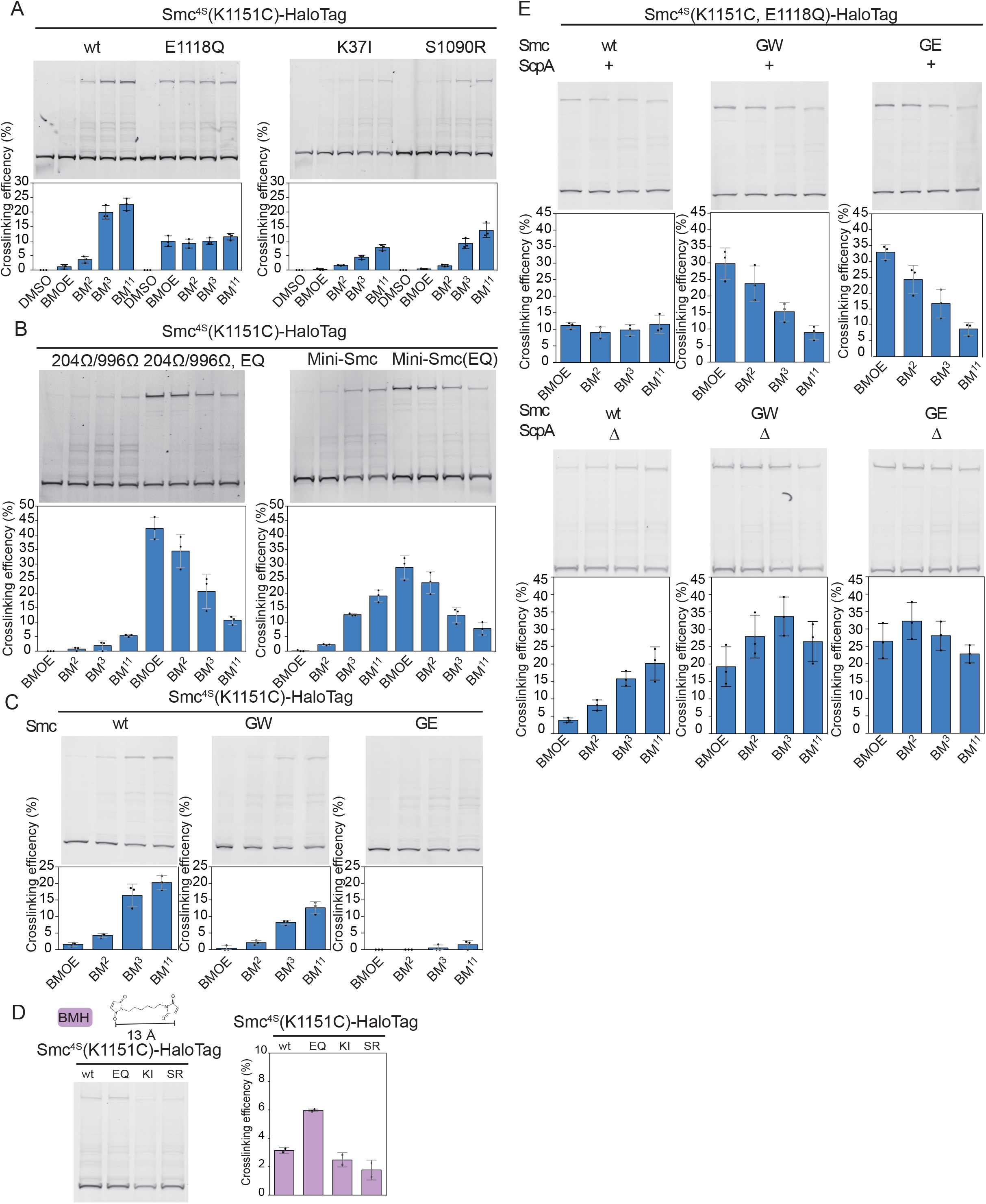
Quantification of *in vivo* cross-linking efficiencies. A. Cross-linking of Smc^4S^(K1151C)-HT and ATPase mutants thereof using different maleimide crosslinkers. Same data as in Figure 5A. B. Cross-linking of Smc^4S^(K1151C)-HT variants harboring arm modifications. Data also shown in Figure 5B. C. Cross-linking of Smc^4S^(K1151C)-HT harboring G302W or G302E mutations. Displayed as Figure 5A. D. Species of Smc^4S^(K1151C)-HT cross-linked with bis-maleimido-hexane (BMH). Displayed as Figure 5A. E. Cross-linking of Smc^4S^(E1118Q, K1151C)-HT variants in the presence (top panels) and absence (bottom panels) of the *scpA* gene. Displayed as Figure 5A.

**Figure S6.**
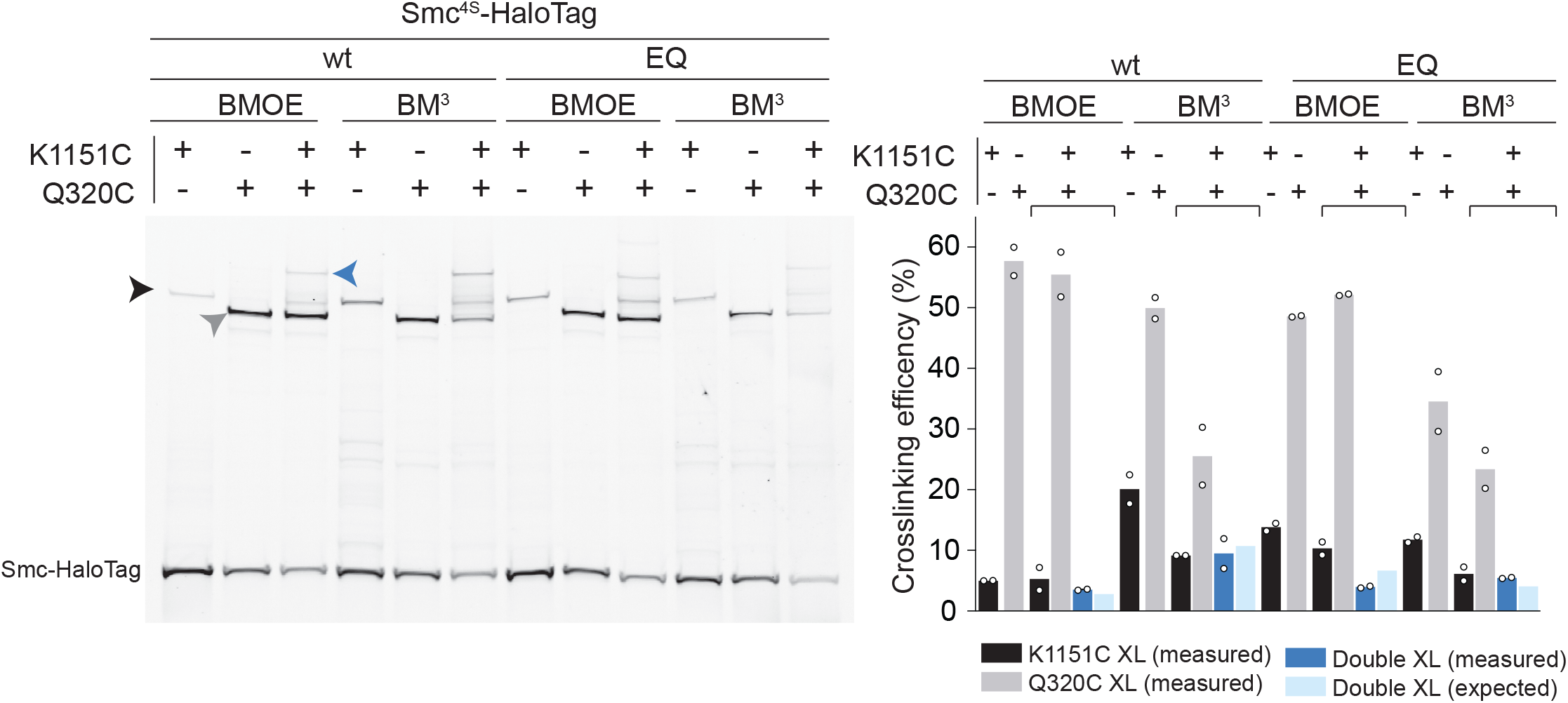
Double cysteine cross-linking *in vivo*. A. BMOE and BM^3^ cross-linking of Smc^4S^-HT and Smc^4S^(EQ)-HT variants harboring single cysteine reporters K1151C or Q320C or a combination of the two cysteines. Quantifications as in Figure 6B.

**Table S1. Related to Figures 4-6.** Strain usage. Genotypes are detailed in Table S5.

| Figure | Strains |
| --- | --- |
| 4C | BSG1002, BSG1007, BSG4097, BSG4099, BSG4095, BSG4098, BSG4096, BSG4094, BSG4092, BSG4093, BSG4037, BSG4038 |
| 4D | BSG1002, BSG1007, BSG4094, BSG4037 |
| 4E | BSG2264, BSG2589, BSG2588, BSG4153, BSG4154, BSG1921, BSG2607, BSG1922, BSG4082, BSG4161 |
| 4F | BSG1002, BSG1007, BSG4037, BSG4094 |
| 4G | BSG1008, BSG1892, BSG4075, BSG4351, BSG4159, BSG4352 |
| 5A | BSG1457, BSG1607, BSG1488 |
| 5B | BSG4160, BSG2693, BSG2531 |
| 6A | BSG1457, BSG4516, BSG4514, BSG1488, BSG4701, BSG4515 |
| S4A | BSG1002, BSG1007, BSG1050, BSG4110, BSG4112, BSG4108, BSG4111, BSG4109, BSG4106, BSG4107 |
| S5A | BSG1457, BSG1488, BSG1607, BSG1600 |
| S5B | BSG2694, BSG2693, BSG2511, BSG2531, BSG1457, BSG4076, BSG4158 |
| S5C | BSG1457, BSG1488, BSG1607, BSG1600 |
| S5D | BSG1488, BSG4077, BSG4160, BSG1509, BSG4168, BSG4170 |
| S6A | BSG1457, BSG1921, BSG4512, BSG1488, BSG1922, BSG4513 |
| S6B | BSG1457, BSG2300, BSG4510, BSG1488, BSG2313, BSG4511 |

**Table S2. Related to Figure 1, S1 and S4.** Mean (µ) and standard deviation (σ) of double gaussian fit to DEER traces. P(r)1 corresponds to the fitted fraction of population 1.

| | $\mu 1$ (nm) | $\sigma 1$ | P(r)1 | $\mu 2$ (nm) | $\sigma 2$ | r.m.s. |
| --- | --- | --- | --- | --- | --- | --- |
| **Smc-ScpAB, apo** |  |  |  |  |  |  |
| D193C, E1118Q | 1.76 | 0.24 | 0.55 | 4.94 | 1.43 | 0.013 |
| D217C, E1118Q | 3.97 | 0.19 | 0.69 | 8.53 | 0.80 | 0.006 |
| R718C, E1118Q | 1.5 | 0.21 | 0.71 | 3.06 | 1.93 | 0.007 |
| R643C, E1118Q | 1.5 | 0.18 | 0.82 | 5.3 | 1.00 | 0.008 |
| D193C | 1.5 | 0.40 | 0.64 | 5.40 | 1.45 | 0.016 |
| D193C, G302W | 1.93 | 0.095 | 0.15 | 5.57 | 1.22 | 0.009 |
| **Smc-ScpAB , ATP, dsDNA<sub>40</sub>** |  |  |  |  |  |  |
| D193C, E1118Q | 1.76 | 0.19 | 0.18 | 5.10 | 1.51 | 0.013 |
| D217C, E1118Q | 3.94 | 0.20 | 0.57 | 8.51 | 1.29 | 0.008 |
| R718C, E1118Q | 1.50 | 0.12 | 0.28 | 2.80 | 2.45 | 0.006 |
| R643C, E1118Q | 1.50 | 0.20 | 0.85 | 5.58 | 0.83 | 0.007 |
| D193C | 1.5 | 0.38 | 0.53 | 4.58 | 2.40 | 0.009 |
| D193C, G302W | 2 (fixed) | 0.69 | N/A | 5.37 | 1.45 | 0.014 |
| **Smc, apo** |  |  |  |  |  |  |
| D193C, E1118Q | 1.70 | 0.22 | 0.53 | 4.85 | 1.50 | 0.016 |
| D217C, E1118Q | 4.03 | 0.23 | 0.66 | 8.66 | 1.09 | 0.007 |
| R718C, E1118Q | 1.5 | 0.19 | 0.74 | 4.01 | 1.56 | 0.007 |
| R643C, E1118Q | 1.5 | 0.17 | 0.83 | 4.92 | 1.21 | 0.007 |
| **Smc, ATP, dsDNA<sub>40</sub>** |  |  |  |  |  |  |
| D193C, E1118Q | 1.66 | 0.20 | 0.18 | 4.60 | 1.86 | 0.016 |
| D217C, E1118Q | 3.99 | 0.25 | 0.44 | 8.45 | 1.74 | 0.009 |
| R718C, E1118Q | 1.5 | 0.20 | 0.20 | 3.95 | 2.16 | 0.005 |
| R643C, E1118Q | 1.5 | 0.17 | 0.83 | 5.03 | 1.21 | 0.007 |

**Table S3. Related to Figure 2.** Crosslinking efficiency of purified Smc^3SV^ single cysteine <u>mutants.</u>

| **Smc3SV** | **Crosslink efficiency (%)** | **SD** |
| --- | --- | --- |
| cysless | N/A | N/A |
| D193C | 85.40 | 0.40 |
| Y944C | 86.47 | 1.57 |
| Q320C | 70.22 | 0.43 |
| A715C | 59.69 | 0.97 |

**Table S4. Related to Figures 2 and 3.** Kinetic parameters of the Smc ATPase.

| **Construct** | **$K_{cat}$ (min<sup>-1</sup>)** | **$K_{0.5}$ (mM)** | **$h$** |
| --- | --- | --- | --- |
| Smc<sup>3SV</sup>(D193C) | 3.9 ± 0.8 | 0.39 ± 0.016 | 1.3 ± 0.3 |
| Smc<sup>3SV</sup>(D193C)-ScpAB | 7.2 ± 0.1 | 0.13 ± 0.007 | 1.1 ± 0.05 |
| Smc<sup>3SV</sup>(D193C) + 3 μM dsDNA<sub>40</sub> | 7.0 ± 0.7 | 0.22 ± 0.05 | 0.9 ± 0.1 |
| Smc<sup>3SV</sup>(D193C)-ScpAB + 3 μM dsDNA<sub>40</sub> | 14.0 ± 0.1 | 0.12 ± 0.003 | 1.3 ± 0.03 |
| Smc<sup>3SV</sup>(D193C) M3M | 0.4 ± 0.4 | 0.36 ± 0.4 | 3.6 ± 10.0 |
| Smc<sup>3SV</sup>(D193C)-ScpAB M3M | 4.7 ± 0.1 | 0.21 ± 0.01 | 1.4 ± 0.08 |
| Smc<sup>3SV</sup>(D193C) M3M + 3 μM dsDNA<sub>40</sub> | 0.7 ± 0.08 | 0.16 ± 0.03 | 2.5 ± 1 |
| Smc<sup>3SV</sup>(D193C)-ScpAB M3M + 3 μM dsDNA<sub>40</sub> | 4.8 ± 0.1 | 0.18 ± 0.01 | 1.4 ± 0.1 |
| Smc<sup>3SV</sup>(E295C) | 6.5 ± 1.4 | 0.13 ± 0.08 | 0.9 ± 0.3 |
| Smc<sup>3SV</sup>(E295C)-ScpAB | 14.4 ± 4.3 | 0.38 ± 0.31 | 0.8 ± 0.2 |
| Smc<sup>3SV</sup>(E295C) + 3 μM dsDNA<sub>40</sub> | 6.8 ± 0.8 | 0.08 ± 0.02 | 1.1 ± 0.3 |
| Smc<sup>3SV</sup>(E295C)-ScpAB + 3 μM dsDNA<sub>40</sub> | 39.2 ± 35.5 | 2.4 ± 5.4 | 0.6 ± 0.1 |
| Smc<sup>3SV</sup>(E295C) M3M | 10.5 ± 0.5 | 0.01 ± 0.007 | 1.5 ± 0.9 |
| Smc<sup>3SV</sup>(E295C)-ScpAB M3M | 30.0 ± 1.4 | 0.14 ± 0.018 | 1.0 ± 0.09 |
| Smc<sup>3SV</sup>(E295C) M3M + 3 μM dsDNA<sub>40</sub> | 11.4 ± 0.4 | 0.02 ± 0.005 | 2.1 ± 1.1 |
| Smc<sup>3SV</sup>(E295C)-ScpAB M3M + 3 μM dsDNA<sub>40</sub> | 29.6 ± 3.2 | 0.14 ± 0.04 | 1.0 ± 0.1 |

## Notes

### Competing Interest Statement

The authors have declared no competing interest.

